# Discovery of cryptic plant diversity in one of the harshest environments: the rooftops of the Alps

**DOI:** 10.1101/2020.06.10.144105

**Authors:** Florian C. Boucher, Cédric Dentant, Sébastien Ibanez, Thibaut Capblancq, Martí Boleda, Louise Boulangeat, Jan Smyčka, Cristina Roquet, Camille Noûs, Sébastien Lavergne

## Abstract

High altitude temperate mountains have long been considered devoid of life owing to high extinction or low speciation rates during the Pleistocene. We performed a phylogenetic and population genomic investigation of an emblematic high-altitude plant clade (*Androsace* sect. *Aretia*, 31 currently recognized species), based on plant surveys conducted during alpinism expeditions. Surprisingly, we inferred that this clade originated in the Miocene and continued diversifying through Pleistocene glaciations, and discovered three novel species of *Androsace* dwelling on different bedrock types on the rooftops of the Alps. This suggests that temperate high mountains have been cradles of plant diversity even during the Pleistocene, with in-situ speciation driven by the combined action of geography and geology. Our findings have an unexpected historical relevance: H.-B. de Saussure likely observed one of these species during his 1788 expedition to the Mont Blanc and we describe it here, over two hundred years after its first sighting.

---

Documenting incipient or recent events of speciation is of paramount importance to understand the mechanisms generating biodiversity, but also to implement adequate conservation actions based on correct species delimitations. This stake is prominent for high-altitude ecosystems, whose biodiversity appears to be disproportionately threatened by ongoing climate change ^1^. In contrast to mountainous environments in general ^2,3^, high-altitude ecosystems are species poor ^4,5^ and are sometimes regarded as ‘the most hostile places on Earth’ (David Attenborough, in 6). The fact that high-altitude environments are biological deserts is generally explained by high extinction rates caused by Pleistocene glaciations ^7^ in combination with low speciation rates caused by low productivity ^8^. In spite of their low diversity however, these environments harbor a high proportion of endemics and retain an important conservation value ^3,4,9^.

In this article we focus on the European Alps (hereafter, ‘the Alps’) and posit the existence of yet undescribed cryptic species within high-altitude plants. This is due to the nature of the two main processes responsible for speciation in this flora: allopatric isolation among mountain ranges and adaptation to divergent substrates ^7,10–12^, which normally leave little imprint on species’ morphology. Several reasons explain why this cryptic diversity might lay unrecognized: (i) Alpine plant taxonomy has mostly been studied using morphology, which may dramatically underestimate diversity ^13^; (ii) phylogenetic studies of Alpine plants have mostly relied on limited sampling within species and limited sequencing effort ^14^, reducing our ability to identify recently diverged lineages; and (iii) high alpine environments have long remained unexplored due to their difficult access, but recent plant surveys suggest that diversity in these environments is much higher than previously assumed ^15^. Improving species delimitation and identification of cryptic species is crucial for a better biodiversity assessment on rooftops of the Alps and to inform conservation strategies for these ecosystems, as done elsewhere and for other organisms ^16,17^. From a broader perspective, accurate species delimitations are also key for testing theories in biogeography ^18,19^ or to enhance our understanding of the speciation process ^20,21^.

Here we study the genus *Androsace* (Primulaceae), which contains some of the vascular plant species dwelling at highest elevations and the coldest places on Earth ^22,23^. We focus on one part of the genus, *Androsace* section *Aretia*, which has its center of diversity in the Alps but is distributed in all European mountains, with some additional species in North America ^24,25^. It has diversified in the last 15 Myr, largely thanks to the emergence of a key innovation, the cushion life form ^26^, that enabled some of its species to conquer the highest altitudes ^25,27^. While understanding the speciation process was our initial incentive to study *Androsace* sect. *Aretia*, we were also intrigued by the recent description of a putative new species in the Mont Blanc range ^28^. A subsequent taxonomic revision then raised the suspicion that other species may remain to be described in this group ^29^, a highly puzzling fact after almost 300 years of study of the Alpine flora since von Haller ^30^.

## Results

We use a large genomic dataset to improve our understanding of the systematics of *Androsace* sect. *Aretia.* We generated ddRAD-seq data ^31^ for 88 individuals spanning the whole distribution range of this clade (Fig. 1a) and 28 out of its 33 currently recognized species (85%), including all European ones (24 spp.). While most species were represented by one to three accessions (see Table S1), we then took advantage of a much denser sampling for a small clade of high-altitude cushion species suspected to include cryptic taxa that was achieved through decade-spanning alpinism expeditions on most higher summits of the Western Alps. The resulting ddRAD tags were aligned to the reference genome of another species from the Primulaceae family, *Primula veris* L. ^32^. This allowed us to infer a high-quality phylogeny of *Androsace* sect. *Aretia* using both maximum-likelihood (hereafter, ‘ML’) on the concatenation of all ddRAD tags (314,363 bp) and species tree inference from unlinked polymorphic sites only (2,461 SNPs). Both methods were largely congruent and led to a highly supported phylogenetic hypothesis (Fig. 1, S2, S3). They supported the split of sect. *Aretia* into two large clades: one of them comprising North-America species as well as mid-elevation specialists from Western Europe (clade /Dicranothrix s*ensu* ^24^), and the second one consisting of high-altitude cushion-forming species from the Alps and adjacent mountain ranges (clade /Eu-Aretia s*ensu* ^24^). Furthermore, ML and species tree inference supported the existence of the same five subclades. One of them, /Douglasia, corresponds to the former genus *Douglasia* Lindley and the four others are named after their most widespread species: /Argentea, /Halleri, /Helvetica and /Vitaliana. All of these clades contain almost exclusively cushion species, except for species of the /Halleri clade, which are perennial rosettes. Divergence time estimation performed using penalized-likelihood on the ML phylogeny using ages derived from a previous study of Primulaceae (*i.e.*, secondary calibration) suggested that all five subclades might have originated before the Pleistocene, possibly in the Pliocene (Fig. 1b).

**Fig. 1.**
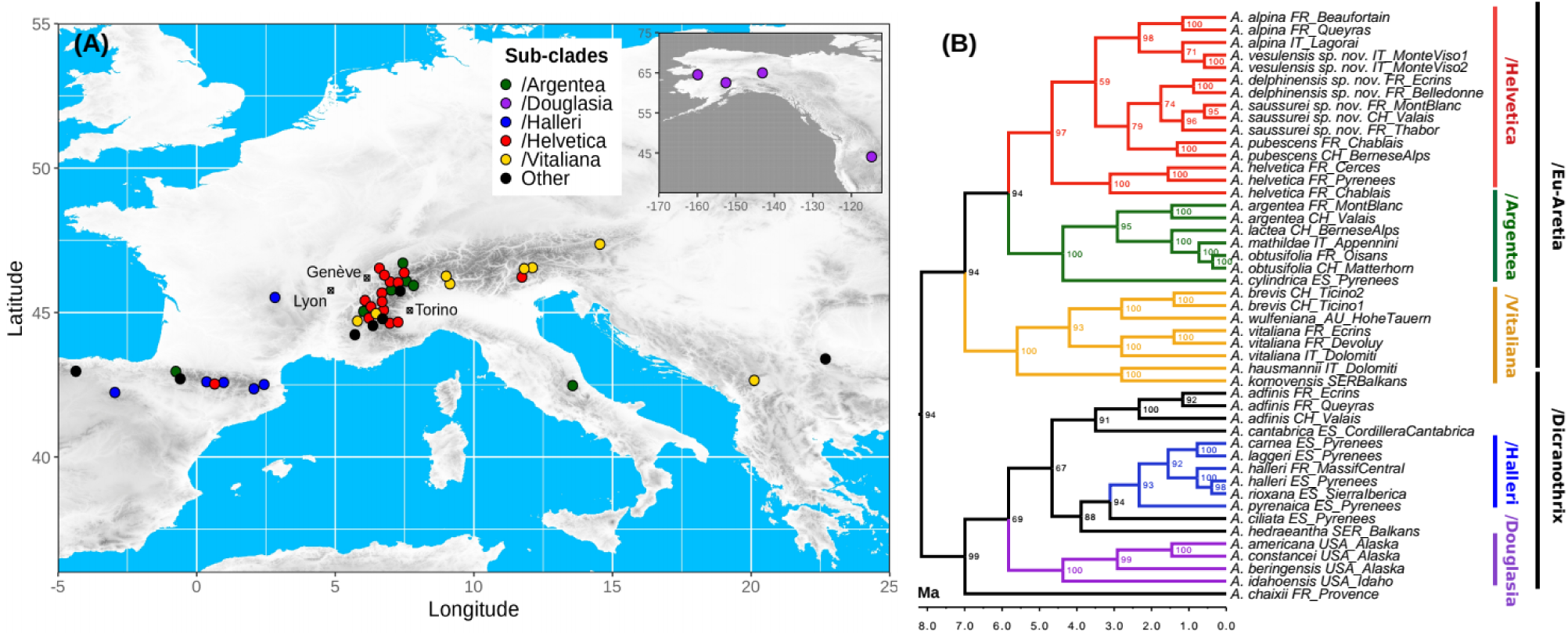
(a) Geographic distribution of samples used for the phylogenomic analysis of *Androsace* sect. *Aretia*. (b) Maximum-likelihood phylogeny of *Androsace* sect. *Aretia* based on the concatenation of 2,700 ddRAD tags together totaling 314,363 bp, and dated using penalized likelihood with secondary calibration. Bootstrap support is displayed at nodes and time is shown on the x axis in million years.

Based on a denser sampling (Fig 2), we then focused on the /Helvetica clade, which was estimated to have originated at 5.8 Ma. This clade contains three currently recognized cushion species growing at high altitudes, *A. alpina, A. helvetica*, and *A. pubescens*, as well as putative new taxa. We first used a constrained clustering method ^33^ to delimit genetic groups without *a priori* taxonomic assignment. Results supported an optimal number of K=7 clusters, which perfectly aligned with the phylogeny of these 51 individuals (Fig. 2). All individuals of *A. helvetica* were assigned to the same cluster. In contrast, individuals morphologically assigned to *A. alpina* were split into two distinct clusters (Fig. 2): one comprising individuals growing on ophiolites of Monte Viso only (hereafter, *A. vesulensis sp. nov.*) and another one comprising all other individuals (hereafter, *A. alpina*). Individuals morphologically assigned to *A. pubescens* were split into three clusters (Fig. 2): one including all individuals growing on limestone (hereafter, *A. pubescens*), one including individuals growing on siliceous substrates in the Mont Blanc and neighboring ranges (hereafter, *A. saussurei sp. nov.*) and the last one including individuals growing on siliceous substrates in the Central French Alps (hereafter, *A. delphinensis sp. nov.*). Finally, further analyses showed that the last cluster (Fig 2), which comprised four individuals with an intriguing morphology, resulted from introgression between *A. pubescens* and *A. saussurei sp. nov.* (see SI).

**Fig. 2.**
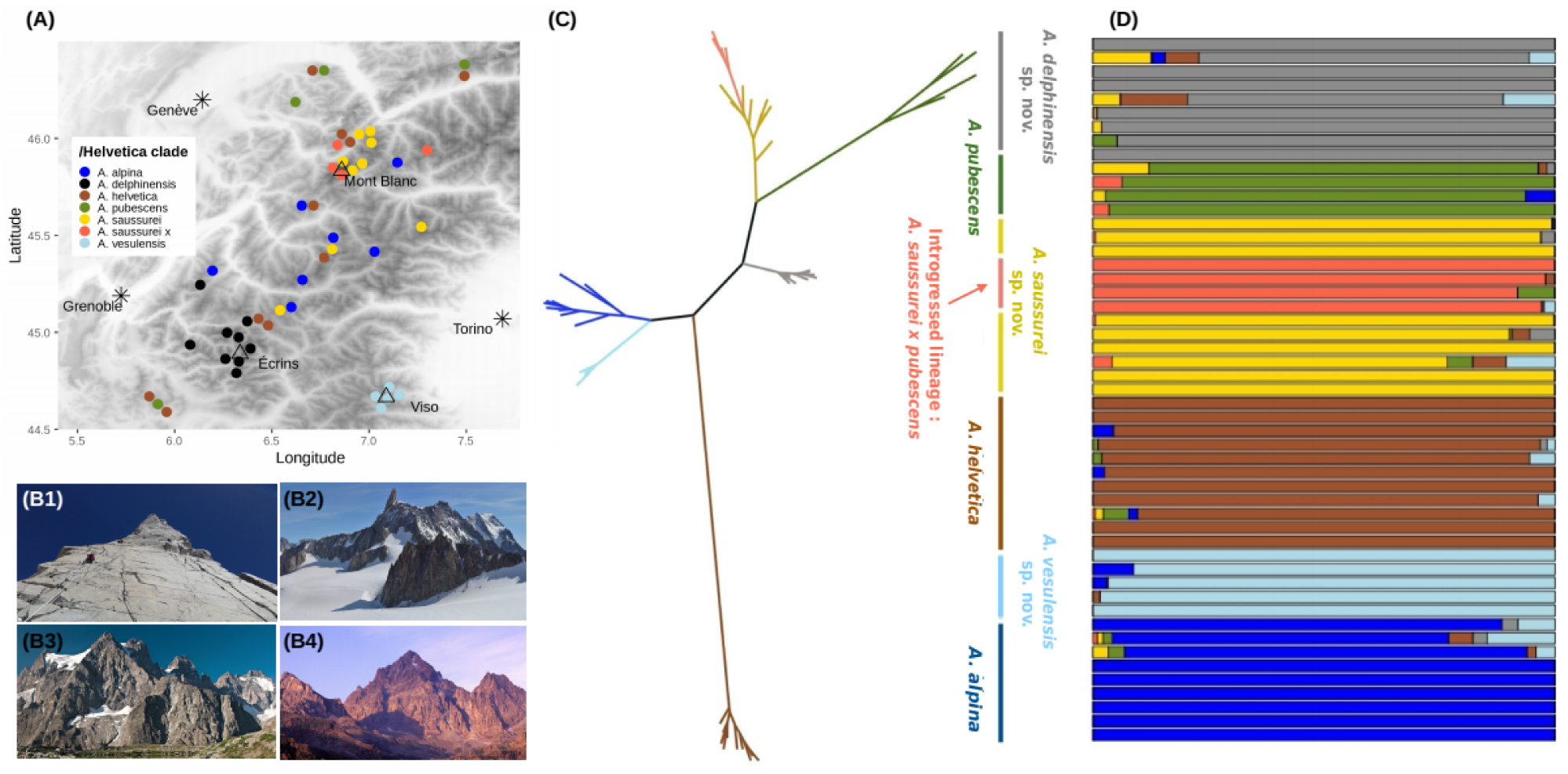
Genetic structure within the /Helvetica clade. (a) Distribution of study samples within the western Alps, spanning the three formerly described species and the three new species described in this study. (b) Pictures depicting the typical high elevation cliff habitats that have been explored for the present study (b1), and the three main mountain ranges where putative novel species occur, namely Mont Blanc (b2), Ecrins (b3), and Monte Viso (b4). (c) Phylogenetic relationships between the 51 individuals of /Helvetica, inferred using ML on a concatenation of 23,780 *loci* (276,745 bp). (d) Assignment of the same individuals to seven genetic clusters as identified based on a strict selection of 381 unlinked SNPs.

The taxonomic status of the genetic clusters that did not align with currently recognized species (*i.e., A. saussurei sp. nov., A. delphinensis sp. nov.* and *A. vesulensis sp. nov.*) was tested using multiple lines of evidence. We first conducted molecular species delimitation, which identifies species as independent evolutionary lineages that do not exchange genes anymore ^34,35^. Bayes Factor Delimitation ^36^ indicated decisive support for a scenario in which *A. alpina* and *A. vesulensis sp. nov.* would be considered different species rather than forming a single one (Bayes Factor, hereafter BF = 1,052 – support for a given scenario is considered decisive when for BF > 150). Using the same approach, we also found decisive support for considering *A. pubescens, A. saussurei sp. nov.* and *A. delphinensis sp. nov.* as distinct species, rather than lumping *A. saussurei sp. nov.* and *A. delphinensis sp. nov.* together (BF = 690), or even lumping the three clusters into a single species, as assumed by current taxonomy (BF = 1,698). Genetic PCAs confirmed that the three new species are distinct genetic groups and not arbitrary portions of a larger genomic cline (see SI). F_ST_ between new species and their close relatives range from 0.08 to 0.3, and are thus of the same order of magnitude as the ones measured between recognized species like *A. alpina* and *A. pubescens* (F_ST_=0.29) or *A. pubescens* and *A. helvetica* (F_ST_=0.17, see SI).

In addition to molecular delimitation, we also used ecological data to confirm the status of these putative species. Based on our extensive prospecting we first refined the chorology and bedrock preferences of these new species and found that they have largely allopatric distributions compared to their close relatives (Fig. 2A, Fig. S15). Contrary to the closely related *A. alpina*, which grows on schist or other siliceous rock screes, *A. vesulensis sp. nov.* grows on more stable rock crevices or cliffs, always on ophiolites. Our new circumscription of *A. pubescens* restricts this species to limestone crevices or cliffs, while the closely related *A. saussurei sp. nov.* and *A. delphinensis sp. nov.* also grow on rock crevices or cliffs, but exclusively on siliceous bedrocks (e.g. granite, quartzite, gneiss, or sandstone). These geographic and edaphic differences thus confirm that the three new taxa deserve species status.

Voucher samples were later examined to look for morphological criteria that could serve for species diagnosis. We found only subtle differences mostly in the morphology of leaves and peduncles trichomes which are described in the taxonomic treatment below (see also Fig. 3b). These morphological difference themselves do not allow distinguishing all six species, but when combined to geographic and edaphic information they can serve to do so (see *Determination key* in SI). Finally, we inferred a species tree for the six newly circumscribed species of /Helvetica using the program SNAPP ^37^. All phylogenetic relationships were strongly supported except for the most basal divergence between *A. helvetica* and the rest of the clade (Fig. 3). Using a prior on the crown age of /Helvetica derived from the dated phylogeny of *Androsace* sect. *Aretia* obtained above, we found that the three newly described species likely originated within the last million years (Fig. S13). These multiple lines of evidence lead us to recognize three new species in *Androsace* sect. *Aretia*: *A. delphinensis, A. saussurei* and *A. vesulensis*, which we typify and formally describe at the end of the manuscript.

**Fig 3.**
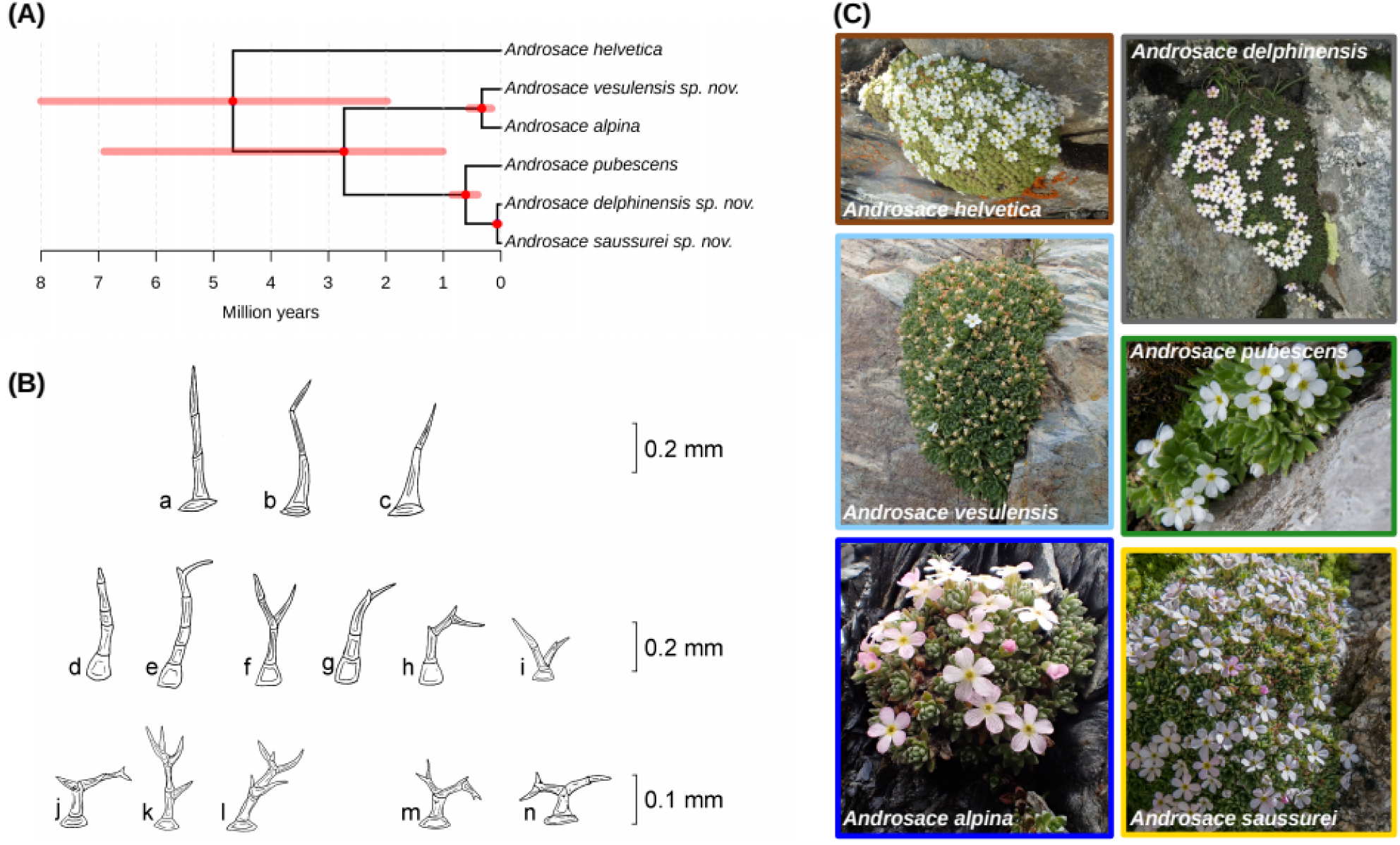
Genetic and morphological delimitations of species within the /Helvetica clade. (A) Species tree inferred from a strict selection of 381 unlinked SNPs, all nodes received a posterior probability of 1.00 except for the most basal one which was supported with 0.45 posterior probability. (B) Drawings of the different trichome morphologies characterizing species of the /Helvetica clade. a-c: *A. pubescens* and *A. helvetica*; d-i: *A. saussurei sp. nov.* and *A. delphinensis sp. nov.* (d-h leaves; f, h, i peduncles); j-l: *A. vesulensis sp. nov.*; m-n: *A. alpina*. All drawings from C. Dentant (C) Pictures depicting the overall morphology of different species delimited within the /Helvetica clade. Pictures from N. Bartalucci, L. Boulangeat, S. Ibanez, and S. Lavergne.

## Discussion

New species are still being frequently described from under-explored regions or under-studied taxonomic groups but it is hardly expected that novel angiosperm species can still be discovered in the European Alpine flora. Against all odds, here we describe three new species of *Androsace* that grow on three of the most emblematic mountain ranges of the Alps: the one that Romans believed was the highest of the World during antiquity, Monte Viso (3841m a.s.l.); one of the wildest ones, the Ecrins (4102m a.s.l.); and the rooftop of Europe, Mont Blanc (4810m a.s.l.). Tribute must be paid to H.-B. de Saussure, who in 1788, during a journey on the Glacier du Géant (3350 m a.s.l, Mont Blanc range), found only one flowering plant species “*sometimes white, sometimes purplish*” ^38^ and made the first observation of what we just described as *Androsace saussurei sp. nov.*, more than two hundred years later. As we describe in alpine *Androsace* here, and as found earlier in arctic species of the genus *Draba* ^13^, cryptic species may have recently arisen within arctico-alpine floras. Our results thus suggest that species diversity may have been underestimated in some high-altitude plant clades, due to limited sequencing and sampling effort of plants dwelling in extreme high alpine environments.

Our study first provides an improved understanding of phylogenetic relationships in *Androsace* sect. *Aretia*. Using thousands of genetic *loci* obtained through ddRAD sequencing along with multiple accessions for the majority of species and mapping these *loci* to a reference genome yields a major improvement in our understanding of the evolutionary history of the whole *Androsace* sect. *Aretia*. It first establishes that geographic structure is marked within the section (Fig. 1), with one clade of North-American species (clade /Douglasia), another one largely confined to the Iberian Peninsula (clade /Halleri), a clade mostly from the Eastern Alps and the Balkans (clade /Vitaliana), and two clades with most of their diversity in the Central and Western Alps (clades /Argentea and /Helvetica). Species bearing the cushion life form, an adaptation to cold and dry environments that has appeared more than a hundred times across angiosperms ^39^, are found in all major subclades of *Androsace* sect. *Aretia* except the one containing species from mid-altitude mountain ranges, /Halleri. Finally, we find that all subclades cited above originated during the Pliocene and continued diversifying throughout the Pleistocene, a period marked by accelerated erosion rates due to global cooling and later glaciations which led to an increase in relief ^40,41^. The formation of deep valleys separating the habitats occupied by species of section *Aretia* might thus have favored its diversification. The precise dating of these events should be taken cautiously since it relies on secondary calibration, but, if anything, these ages are overestimated given that the original study had obtained rather old divergence times compared to other studies of Primulaceae ^7^.

Our study provides evidence for recent plant species origination in some of the most extreme environments on Earth. High-altitude habitats in the Alps host relatively little plant life ^4,22^ and have thus been long viewed as historical sinks of diversity, due to low speciation owing to harsh climatic conditions and to high extinction driven by Pleistocene glaciations. Instead, we show that the number of species recognized in the /Helvetica clade should be raised from three to six, and that three speciation events likely took place during the Pleistocene, a period during which the Alps were largely covered by glaciers. The occurrence of Pleistocene speciation, together with the distribution of species in the interior of the Alps (Fig. 1), suggests that plants of the /Helvetica clade may have survived in nunataks located above glaciers and even diversified during glacial periods ^42^. This is a plausible scenario given that these plants frequently grow on high-altitude cliffs above glaciers today. For species of /Helvetica it has even been suggested that these refugia might have been located both in the center and in the periphery of the Alps ^43^. More generally, the origin of major subclades within *Androsace* sect. *Aretia* in the Pliocene and the occurrence of several speciation events during the Pleistocene supports the idea that mountain floras worldwide are the result of fast and recent evolutionary radiations ^44^.

The new species described in this study have allopatric distributions with their closest relatives (Fig. 2). This supports the prominent role of allopatric speciation in the flora of the European Alpine System, which can be conceived as an archipelago of ‘sky islands’ among which gene flow is probably very limited ^7,11,45^. But the other striking observation is that these new species grow on distinct substrates compared to their close relatives. While *A. pubescens* exclusively grows on limestone, its sister clade, which is formed of *A. saussurei sp. nov.* and *A. delphinensis sp. nov.* is found on siliceous rocks. Similarly, the widespread *A. alpina* is restricted to siliceous rocks whereas its sister species *A. vesulensis* grows on mafic or ultramafic ophiolites. Substrate is known to have played an important role in the phylogeography of the Alpine flora ^46^ and has also been proposed as a driver of speciation ^7,12,47^. This is likely because of the trade-offs required for adaptation to alternative soil chemistries, but also because bedrock type is typically uniform across spatial scales of a few kilometers in the Alps and because it has remained constant during the whole geological history of the Alps, except for the erosion of some sedimentary rocks capping nowadays exposed igneous ones ^48^. This illustrates that allopatric and ecological speciation are intertwined rather than mutually-exclusive mechanisms. In contrast, climate doesn’t seem to be an axis of niche divergence in the Alpine flora ^7^, probably because it varies over short distances along elevational gradients and because it has varied drastically over timescales of a few thousand years following glacial cycles. In /Helvetica, the spatial separation and temporal constancy of bedrock types would have triggered ecological speciation, leaving enough time for different plant lineages to adapt to different substrates, while the same may not have been possible for climate. Plants of the /Helvetica clade would be especially prone to substrate specialization given that they live on cracks or screes of the bedrock and are thus directly affected by it’s properties. Importantly, reproductive isolation does not seem to be complete between species, as exemplified by the existence of hybrids between *A. pubescens* and *A. helvetica* ^*43*^ or *A. saussurei sp. nov.* as we have found here. This could be the result of allopatric divergence, during which intrinsic reproductive barriers are not selected and thus evolve rather slowly ^49^.

This systematic study of one of the plant clades found in the coldest environments on Earth bears importance for implementing appropriate conservation strategies in ecosystems at stake with the issue of climate change. We just unraveled the existence of rather cryptic species of high alpine *Androsace* using molecular data, but further scrutiny of fine morphological characters allowed proposing diagnostic traits for these new species. Despite the seemingly good news of this finding, the newly identified *A. delphinensis sp. nov., A. saussurei sp. nov.*, and *A. vesulensis sp. nov.* are already facing important threats. Until a proper threat assessment is performed, these novel species inherit the protection status of the species they were previously included in (that is, *A. pubescens* and *A. alpina*) and are thus automatically protected in France, Italy, and Switzerland. But, given their restricted range sizes (Fig. 2) and the important risk they face from climate change, either directly ^1^ or due to increased competition with colonizers from lower altitudes ^50^, there is little doubt that these novel species will require species-specific protection plans. Here we have only investigated the systematics of one genus, but high-altitude ecosystems probably harbor many more unknown species that may disappear before ever being described. It is thus time to change our perception of these ecosystems, which are much more deserts of knowledge than deserts of life.

## Methods

Samples used in this study were collected during a number of botanical expeditions spanning large elevation gradients on over 90 highest summits and passes of the Western Alps, most of the time requiring the means of alpinism techniques. For most of these high elevation sites, no botanical data had ever been recorded. This study used 88 individuals sampled throughout European mountains (Fig. 1a), including 1-3 individuals of all 24 European species of *Androsace* sect. *Aretia*, and a denser coverage of *A. alpina* (L.) Lam. (14 individuals, including one suspected new taxon), *A. helvetica* (L.) All. (11 individuals), and *A. pubescens* DC. (26 individuals, including two suspected new taxa). We also included one sample of four of the nine North-American species of the section, plus three outgroups (Table S1).

Total DNA from all samples was extracted using a DNeasy Plant Mini Kit (Quiagen, Germany). A double-digested RAD (restriction site associated DNA) experiment was conducted using a modified version of the original protocol (^31^, see SI), generating more than 275 million reads of 2 x 125bp. Rather than relying on *de novo* assembly of these loci, we preferred to align them on the reference genome of another species from the Primulaceae family: *Primula veris* L. ^32^. We also checked that technical replicates of our various ddRAD-seq libraries and sequencing runs produced comparable data prior to combining them for further analyses (Fig. S1).

Phylogenetic relationships within *Androsace* sect. *Aretia* were inferred using two different approaches: ML inference of all ddRADseq tags concatenated (314,363 bp) using IQ-TREE ^51^ and species tree inference from 2,461 unlinked SNPs using SVDquartets ^52^. In both cases, clade support was measured using bootstrap and trees were rooted thanks to the inclusion of three outgroup taxa from the Primulaceae family, located at increasing phylogenetic distances from our ingroup: *A. septentrionalis* L., *Primula hirsuta* All., and *Lysimachia nummularia* L. (see SI). We then dated the phylogram obtained from concatenation using penalized likelihood ^53^. We calibrated two nodes of the phylogeny using median ages estimates from a previous study of Primulaceae ^7^: the crown node of *Androsace* sect. *Aretia* (8.16 Ma) and the divergence between the genera *Androsace* and *Primula* (33.3 Ma).

In order to revise the systematics of /Helvetica and to test the putative species status of newly discovered taxa, we used an integrative taxonomic approach combining genomics, geography, and habitat characterization.We started by doing a different SNP calling for the 51 individuals of the /Helvetica clade using the same pipeline as described above, which resulted in 23,780 loci aligned to the reference genome, containing a total of 276,745 bp and 7,806 SNPs. In order to use population genetic measures on this recently diverged clade, we strictly filtered this initial SNP dataset: we only kept biallelic SNPs that had less than 40% missing data, that had minor allele frequencies >4%, and that were unlinked, yielding a final dataset of 381 SNPs. This number is rather low but we preferred working with these high-quality SNPs only rather than relying on *de novo* assembly of ddRADseq tags, which gave similar results (see SI). Genetic clusters were then inferred with no *a priori* using an algorithm aimed at estimating individual ancestry ^33^. We tested numbers of genetic clusters K ranging from 1 to 10 and the optimal number of clusters was chosen based on the cross-entropy of the best run for each value of K ^33^. We tested the species status of the five clusters that did not correspond to already recognized species using Bayes factor delimitation ^36^. This method relies on comparing the marginal likelihood of alternative species delimitation scenarios using under the multispecies coalescent model to identify evolutionary lineages that do not exchange genes anymore, *i.e.* species under the general species concept ^34,35^. For each genetic cluster, alternative species delimitation scenarios in which the cluster would have species rank or not were statistically compared (Fig. S10). The geographic distribution and bedrock affinities of all species were determined thanks to the field data collected during our own sampling campaigns, in combination to data obtained from an online citizen-science project dedicated to the digitization and geo-referencing of herbarium records of the genus *Androsace* (http://lesherbonautes.mnhn.fr/missions/13798338). We also scrutinized 139 plant samples and coded a number of morphological characters, in order to provide morphological diagnosis criteria to differentiate study taxa. Once we had confirmed that these six taxa deserved species status, we inferred their phylogenetic relationships using species tree inference from unlinked SNPs ^37^. Four independent chains of 500,000 steps were combined to produce a maximum clade credibility tree, which was calibrated to absolute time using the estimation of the crown age of /Helvetica obtained above.

## Taxonomic treatment: description of three novel *Androsace* species

### *\*Androsace delphinensis* Dentant, Lavergne, F.C. Boucher & S. Ibanez sp. nov

[=*Androsace pubescens* auct. non DC.]

Holotypus (designed here): France, Hautes-Alpes, Ecrins, Pic Coolidge (3775m a.s.l.), close to the summit, in south-facing crevices in granite, [6,358341°N/44,909247°], 3740m a.s.l., 2019/08/13. Coll.: Dentant. G[G00419022]! – Isotypus: GRM[MHNGr.2020.46703]!

Perennial plant forming a creeping cushion, sometimes compact, up to 5(10) cm high, 3–20 cm in diameter, made of loose to slightly compact rosettes. Leaves lanceolate, 7–10 × 1-1.5 mm, hairy on both sides. **Hairs** persistent, **simple** (proportion (0)25% to 75(100)%) **bifurcated** or **branched** (at least 3 branches), slightly curved, 0.2–0.5 mm long, composed of (3)4-6(8) cells. Hairs often branched with a short branch at the top, sometimes broken. **Pedicel with hairs exclusively bifurcated or branched. Corolla always white**, 7-8 mm in diameter. Flowering: June to August. Substratum: gneiss, granite, sandstone, flysch. Altitude: 2400m to 3850m a.s.l.. Chorology: Southwestern Alps.

### *\*Androsace vesulensis* Dentant, Lavergne, F.C. Boucher & S. Ibanez sp. nov

[=*Androsace pubescens* auct. non DC.; =*Androsace alpina* auct. non Lam.]

Holotypus (designed here): Italy, Piedmont, Monte Viso (3841m a.s.l.), from passo Giacoletti, on the north ridge of Punta Gastaldi, in crevices of ophiolite, [7,0758520°/44,6842880°], 3020m a.s.l., 2017/08/04. Coll.: Lavergne & Ibanez. G[G00419023]! – Isotypus: GRM[MHNGr.2020.46704]!

Perennial plant forming a creeping cushion, sometimes compact, up to 5(8) cm high, 3–10 cm in diameter, made of loose to slightly compact rosettes. Leaves lanceolate, 5–6.3 × 1–2.2 mm, hairy (mainly on edges). Hairs persistent, **deer-antler-shaped**, 0.1–0.2 mm long, composed of 3–4(7) cells. **Corolla always white**, 7 mm in diameter. Flowering: June to August. Substratum: ophiolite (basalt, gabbro, serpentine). Altitude: 2800m to 3800m a.s.l. Chorology: endemic of Monte Viso and neighboring summits (Italy and France).

### *\*Androsace saussurei* Dentant, Lavergne, F.C. Boucher & S. Ibanez sp. nov

[=*Androsace pubescens* auct. non DC.; =*Androsace alpina* auct. non Lam.]

Holotypus (designed here): France, Haute-Savoie, Chamonix, Western ridge of the Aiguille du Passon, from the col du Passon, in crevices of granite [6,986936°/45,976718°], 3170m a.s.l., 2019/07/17, coll.: Lavergne & Ibanez. G[G00419024]! – Isotypus: GRM[MHNGr.2020.46705]!

Perennial plant forming a creeping cushion, sometimes compact, up to 5(8) cm high, 3–10 cm in diameter, made of loose to slightly compact rosettes, 4.7-8.1mm in diameter. Leaves lanceolate, 4.5–6(9.5) × 1.1–1.7 mm, hairy on both surfaces, **often reddish at the tips** (higher anthocyanin concentration). **Hairs** persistent, **simple** (proportion (0)25% to 75(100)%), **bifurcated** or **branched** (at least 3 branches), 0.2–0.5 mm long, composed of 3–5(8) cells. Pedicel with hairs exclusively bifurcated or branched. **Corolla white to purplish**, 7 mm in diameter. **Buds often purplish**. Flowering: June to August. Substratum: protogine, granite. Altitude: 2000m to 4060m a.s.l. – (highest elevation known for a vascular plant in Italy, observed by one of us (S. Ibanez) in the south side of Mont Blanc). Chorology: Western Alps.

## Acknowledgements

We thank all people who contributed to sampling: N. Bartalucci, T. Bulle, J. Charron, R. Douzet, P. Dufour, S. Eggenberg, L. Gallien, L. Garraud, E. Hustache, S. Julien, M. Kolářová, P. Koutecký, S. Latzin, A. Moehl, J.Pilátová, D. Požárová, P. Saccone, P. Schönswetter, G. Schneeweiss, M. Smyčková, J. Van Es, J.-C. Villaret, and S. Wipf. P. Saccone and C.H. Albert participated in the early stage of this project. The ALA and ID herbaria kindly provided material for North American species and J. Renaud helped with designing the sampling. The research was funded by the ANR project Origin-Alps (ANR-16-CE93-0004). Some samples of this study were collected as part of the PhyloAlps project. The sampling campaign in the Balkans region was supported by Bourse de Terrain of the French Ecological Society to J. Smyčka. This study was performed with all necessary authorizations from the following French departments, Savoie (n°2017-985), Haute Savoie (n°2017-1423), Hautes-Alpes (n°<u>05-2017-07-21-001</u>), Isère (n°<u>38-2017-07-20-003</u>), Alpes Maritimes (n°<u>06-2017-07-18</u>), along with relevant research authorizations from protected natural areas. The authors are extremely grateful to the many virtual botanists that have contributed to the *Herbonautes* citizen science project called <u>“Les Androsaces, des primulacées d’altitude”</u>, and managed by the National Museum of Natural History, Paris.

## References

1. Engler, R. et al. 21st century climate change threatens mountain flora unequally across Europe. Glob. Chang. Biol. 17, 2330–2341 (2011).

2. Spehn, E. M. & Körner, C. A global assessment of mountain biodiversity and its function. Glob. Chang. Mt. Reg. 393–400 (2005).

3. Rahbek, C. et al. Humboldt’s enigma: What causes global patterns of mountain biodiversity? Science (80-.). 365, 1108–1113 (2019).

4. Aeschimann, D., Rasolofo, N. & Theurillat, J. Analyse de la flore des Alpes. 3: biologie et phénologie. Candollea 67, 3–22 (2012).

5. McCain, C. M. & Grytnes, J.-A. Elevational Gradients in Species Richness. Encyclopedia of Life Sciences (2010).

6. Anderson, J. Planet Earth II, Episode 2: Mountains. (British Broadcasting Corporation, 2016).

7. Boucher, F. C., Zimmermann, N. E. & Conti, E. Allopatric speciation with little niche divergence is common among alpine Primulaceae. J. Biogeogr. 43, 591–602 (2016).

8. Körner, C. Alpine Plant Life. (Springer, 1999).

9. Smyčka, J. et al. Disentangling drivers of plant endemism and diversification in the European Alps – A phylogenetic and spatially explicit approach. Perspect. Plant Ecol. Evol. Syst. 28, 19–27 (2017).

10. Conti, E., Soltis, D. E., Hardig, T. M. & Schneider, J. Phylogenetic relationships of the silver saxifrages (Saxifraga, sect. Ligulatae haworth): implications for the evolution of substrate specificity, life histories, and biogeography. Mol. Phylogenet. Evol. 13, 536–555 (1999).

11. Kadereit, J. W., Griebeler, E. M. & Comes, H. P. Quaternary diversification in European alpine plants: pattern and process. Philos. Trans. R. Soc. Lond. B. Biol. Sci. 359, 265–274 (2004).

12. Moore, A. J. & Kadereit, J. W. The evolution of substrate differentiation in Minuartia series Laricifoliae (Caryophyllaceae) in the European Alps: In situ origin or repeated colonization? Am. J. Bot. 100, 2412–2425 (2013).

13. Grundt, H. H., Kjølner, S., Borgen, L., Rieseberg, L. H. & Brochmann, C. High biological species diversity in the arctic flora. Proc. Natl. Acad. Sci. U. S. A. 103, 972–975 (2006).

14. Kadereit, J. W. The role of in situ species diversification for the evolution of high vascular plant species diversity in the European Alps—A review and interpretation of phylogenetic studies of the endemic flora of the Alps. Perspect. Plant Ecol. Evol. Syst. 26, 28–38 (2017).

15. Marx, H. E. et al. Riders in the sky (islands): Using a mega-phylogenetic approach to understand plant species distribution and coexistence at the altitudinal limits of angiosperm plant life. J. Biogeogr. 44, 2618–2630 (2017).

16. Hebert, P. D. N., Penton, E. H., Burns, J. M., Janzen, D. H. & Hallwachs, W. Ten species in one: DNA barcoding reveals cryptic species in the neotropical skipper butterfly Astraptes fulgerator. Proc. Natl. Acad. Sci. U. S. A. 101, 14812 LP – 14817 (2004).

17. Vieites, D. R. et al. Vast underestimation of Madagascar’s biodiversity evidenced by an integrative amphibian inventory. Proc. Natl. Acad. Sci. 106, 8267 LP – 8272 (2009).

18. Bálint, M. et al. Cryptic biodiversity loss linked to global climate change. Nat. Clim. Chang. 1, 313–318 (2011).

19. Gill, B. A. et al. Cryptic species diversity reveals biogeographic support for the ‘mountain passes are higher in the tropics’ hypothesis. Proc. R. Soc. B Biol. Sci. 283, 20160553 (2016).

20. Moritz, C. et al. Identification and dynamics of a cryptic suture zone in tropical rainforest. Proc. R. Soc. B Biol. Sci. 276, 1235–1244 (2009).

21. Struck, T. H. et al. Finding Evolutionary Processes Hidden in Cryptic Species. Trends Ecol. Evol. 33, 153–163 (2018).

22. Körner, C. Coldest places on earth with angiosperm plant life. Alp. Bot. 121, 11–22 (2011).

23. Dentant, C. The highest vascular plants on Earth. Alp. Bot. (2018). doi: 10.1007/s00035-018-0208-3

24. Schneeweiss, G., Schönswetter, P., Kelso, S. & Niklfeld, H. Complex biogeographic patterns in Androsace (Primulaceae) and related genera: evidence from phylogenetic analyses of nuclear internal transcribed spacer and plastid trnL-F sequences. Syst. Biol. 53, 856–876 (2004).

25. Boucher, F. C. et al. Reconstructing the origins of high-alpine niches and cushion life form in the genus Androsace s.l. (Primulaceae). Evolution (N. Y). 66, 1255–1268 (2012).

26. Aubert, S., Boucher, F., Lavergne, S., Renaud, J. & Choler, P. 1914-2014: A revised worldwide catalogue of cushion plants 100 years after Hauri and Schröter. Alp. Bot. 124, 59–70 (2014).

27. Roquet, C., Boucher, F. C., Thuiller, W. & Lavergne, S. Replicated radiations of the alpine genus Androsace (Primulaceae) driven by range expansion and convergent key innovations. J. Biogeogr. 40, 1874–1886 (2013).

28. Jordan, D. La flore rare ou menacée de Haute-Savoie. (Naturalia publications, 2015).

29. Dentant, C., Lavergne, S. & Malécot, V. Taxonomic revision of West-Alpine cushion plant species belonging to Androsace subsect. Aretia. Bot. Lett. 165, 337–351 (2018).

30. von Haller, A. Enumeratio methodica stirpium Helvetiae indigenarum. (1742).

31. Peterson, B. K., Weber, J. N., Kay, E. H., Fisher, H. S. & Hoekstra, H. E. Double Digest RADseq: An Inexpensive Method for De Novo SNP Discovery and Genotyping in Model and Non-Model Species. PLoS One 7, e37135 (2012).

32. Nowak, M. D. et al. The draft genome of Primula veris yields insights into the molecular basis of heterostyly. Genome Biol. 16, 12 (2015).

33. Frichot, E., Mathieu, F., Trouillon, T., Bouchard, G. & François, O. Fast and Efficient Estimation of Individual Ancestry Coefficients. Genetics 196, 973 LP – 983 (2014).

34. De Queiroz, K. Species Concepts and Species Delimitation. Syst. Biol. 56, 879–886 (2007).

35. Fujita, M. K., Leaché, A. D., Burbrink, F. T., McGuire, J. A. & Moritz, C. Coalescent-based species delimitation in an integrative taxonomy. Trends Ecol. Evol. 27, 480–488 (2012).

36. Leaché, A. D., Fujita, M. K., Minin, V. N. & Bouckaert, R. R. Species delimitation using genome-wide SNP data. Syst. Biol. 63, 534–542 (2014).

37. Bryant, D., Bouckaert, R., Felsenstein, J., Rosenberg, N. A. & RoyChoudhury, A. Inferring species trees directly from biallelic genetic markers: bypassing gene trees in a full coalescent analysis. Mol. Biol. Evol. 29, 1917–1932 (2012).

38. de Saussure, H.-B. Voyage dans les Alpes - tomes I, II, III et IV. (Samuel Fauche (tome I); Barde, Manget & Compagnie (tome II); Louis Fauche-Borel (tomes III&IV)).

39. Boucher, F. C., Lavergne, S., Basile, M., Choler, P. & Aubert, S. Evolution and biogeography of the cushion life form in angiosperms. Perspect. Plant Ecol. Evol. Syst. 20, 22–31 (2016).

40. Valla, P. G., Shuster, D. L. & Van Der Beek, P. A. Significant increase in relief of the European Alps during mid-Pleistocene glaciations. Nat. Geosci. 4, 688–692 (2011).

41. Herman, F. et al. Worldwide acceleration of mountain erosion under a cooling climate. Nature 504, 423–426 (2013).

42. Schneeweiss, G. M. & Schönswetter, P. A re-appraisal of nunatak survival in arctic-alpine phylogeography. Mol. Ecol. 20, 190–192 (2011).

43. Schneeweiss, G. M., Winkler, M. & Schönswetter, P. Secondary contact after divergence in allopatry explains current lack of ecogeographical isolation in two hybridizing alpine plant species. J. Biogeogr. 44, 2575–2584 (2017).

44. Hughes, C. E. & Atchison, G. W. The ubiquity of alpine plant radiations: from the Andes to the Hengduan Mountains. New Phytol. 207, 275–282 (2015).

45. Ozenda, P. L’endémisme au niveau de l’ensemble du Système alpin. Acta Bot. Gall. 142, 753–762 (1995).

46. Alvarez, N. et al. History or ecology? Substrate type as a major driver of spatial genetic structure in Alpine plants. Ecol. Lett. 12, 632–640 (2009).

47. Conti, E., Soltis, D. E., Hardig, T. M. & Schneider, J. Phylogenetic Relationships of the Silver Saxifrages (Saxifraga, Sect. Ligulatae Haworth): Implications for the Evolution of Substrate Specificity, Life Histories, and Biogeography. 13, 536–555 (1999).

48. Glotzbach, C., Beek, P. & Spiegel, C. Episodic exhumation and relief growth in the Mont Blanc massif, Western Alps from numerical modelling of thermochronology data. Earth Planet. Sci. Lett. 304, 417–430 (2011).

49. Nosil, P. Ecological speciation. (Oxford University Press, 2012).

50. Steinbauer, M. J. et al. Accelerated increase in plant species richness on mountain summits is linked to warming. Nature 556, 231–234 (2018).

51. Nguyen, L.-T., Schmidt, H. A., von Haeseler, A. & Minh, B. Q. IQ-TREE: a fast and effective stochastic algorithm for estimating maximum-likelihood phylogenies. Mol. Biol. Evol. 32, 268–274 (2015).

52. Chifman, J. & Kubatko, L. Quartet Inference from SNP Data Under the Coalescent Model. Bioinformatics 30, 3317–3324 (2014).

53. Sanderson, M. J. Estimating Absolute Rates of Molecular Evolution and Divergence Times: A Penalized Likelihood Approach. Mol. Biol. Evol. 19, 101–109 (2002).

