## supplementary information for "Discovery of cryptic plant diversity in one of the harshest environments: the rooftops of the Alps"

| Sample code | Species name | Clade | Country | Mountain range / Region | Locality | Date (d/m/y) | Elevation | Lat | Long | Collectors | Raw number of reads | Number of reads after quality filtering |
|---|---|---|---|---|---|---|---|---|---|---|---|---|
| **A_ADF_ADF** | *A. adfinis* Biroli | na | France | Ecrins | Serre Eyraud | na | na | 44,653165 | 6,311374 | C. Dentant | 2837087 | 2836687 |
| **A_ADF_BRI** | *A. adfinis* Biroli | na | France | Queyras | Col de l'Izoard | na | na | 44,819775394136 | 6,73513320166106 | R. Douzet | 2781539 | 2781130 |
| **A_ADF_PUB** | *A. adfinis* Biroli | na | Italy | Valais | Col du Grand Saint Bernard | 11/06/2010 | na | 45,8708953677333 | 7,27344809694678 | A. Moehl | 3380353 | 3379900 |
| **AA11** | *A. alpina* (L.) Lam | /Helvetica | France | Vanoise | Plan Richard | 14/08/2016 | 2605 | 45,4876516 | 6,8163767 | S. Lavergne | 2561222 | 2560802 |
| **AA22** | *A. alpina* (L.) Lam | /Helvetica | France | Thabor | Tour du Cheval Blanc, scree slopes below the S face | 27/06/2017 | 2686 | 45,124838 | 6,581129 | S. Lavergne, M. Boleda | 3370379 | 3369890 |
| **AA26** | *A. alpina* (L.) Lam | /Helvetica | France | Beaufortain | Col de la Nova | 30/06/2017 | 2770 | 45,653549 | 6,683854 | S. Lavergne, P. Dufour | 7097905 | 7096889 |
| **AA28** | *A. alpina* (L.) Lam | /Helvetica | Italy | Lagorai | Colbricon | 09/07/2017 | 2516 | 46,27461 | 11,75194 | S. Lavergne, J. Smycka | 4142591 | 4141986 |
| **AA30** | *A. alpina* (L.) Lam | /Helvetica | Italy | Bergamo Alps | Cima del Desenigo, 100m of S summit | 11/07/2017 | 2760 | 46,20019 | 9,56335 | S. Lavergne, J. Smycka | 3587079 | 3586499 |
| **AA4** | *A. alpina* (L.) Lam | /Helvetica | France | Vanoise | Col de Chavière | 24/07/2016 | 2796 | 45,270685 | 6,657427 | L. Boulangeat | 3834089 | 3833514 |
| **AL1** | *A. alpina* (L.) Lams | /Helvetica | Italy | Valais | Fenetre de Ferret | 02/08/2018 | 2688 | 45,87556667 | 7,14555 | S. Lavergne, F. Boucher, L. Boulangeat | 1006979 | 1006840 |
| **AL2** | *A. alpina* (L.) Lam | /Helvetica | France | Belledonne | Col de Villonet | 14/07/2018 | 2460 | 45,3179 | 6,1946 | F. Boucher | 1524197 | 1524032 |
| **AL3** | *A. alpina* (L.) Lam | /Helvetica | France | Vanoise | Col de l'Iseran | 11/07/2018 | 2790 | 45,4158 | 7,0282 | F. Boucher | 2175159 | 2174881 |
| **D_ARC** | *A. americana* Wendelbo | /Douglasia | USA | Alaska | Alaska (herbarium specimen ALA) | 08/01/2001 | na | 62,53283333 | -152,571 | na | 622839 | 622766 |
| **AV_217** | *A. argentea* (C.F.Gaertn.) Lapeyr. | /Argentea | France | Mont Blanc | Minaret | 09/2017 | 3200 | 45,9532417 | 7,014275 | S. Ibanez | 4829781 | 4829182 |
| **AV_226** | *A. argentea* (C.F.Gaertn.) Lapeyr. | /Argentea | Switzerland | Valais | Dent Blanche | 07/2017 | 3469 | 46,033 | 7,6 | S. Ibanez | 4213054 | 4212504 |
| **D_BER** | A. beringensis (S.Kelso, Jurtzev & D.F.Murray) Cubey | /Douglasia | USA | Alaska | Alaska (herbarium specimen ALA) | 29/06/1998 | na | 64,51666667 | -159,88333333 | na | 766922 | 766819 |
| **A_BRE_1** | *A. brevis* (Hegetschw.) Ces. | /Vitaliana | Switzerland | Ticino | Piano Cuescio | 07/07/2010 | na | 46,1179665113497 | 9,07250666500613 | S. Wipf | 2159337 | 2159045 |
| **A_BRE_2** | *A. brevis* (Hegetschw.) Ces. | /Vitaliana | Switzerland | Ticino | Piano Cuescio | 07/07/2010 | na | 46,1347355277287 | 9,06394959163893 | S. Wipf | 3095856 | 3095451 |
| **A_CAN** | *A. cantabrica* (Losa & P.Monts.) Kress | na | Spain | Cordillera Cantabrica | Sestil Alto | 23/07/2016 | 2075 | 42,967 | -4,3542222 | S. Lavergne, C. Roquet, J. Smycka | 1652891 | 1652696 |
| **A_CAR_ROS** | *A. carnea* L. | /Halleri | Spain | Pyrenees | Living plant collection of the Lautaret Alpine Garden | 29/07/2015 | na | na | na | S. Lavergne, M. Boleda | 1299514 | 1299350 |
| **A_CHA** | *A. chaixii* Gren. & Godr. | na | France | Baronnies | Combe Marsenne | 05/05/2009 | na | 44,2325939949821 | 5,70907000510294 | J. Van Es | 2689684 | 2689375 |
| **A_CIL** | *A. ciliata* DC. | na | Spain | Pyrenees | Bisaurin | 08/07/1905 | 2580 | 42,7877025 | -0,6431169 | C. Roquet | 3218241 | 3217847 |
| **D_GOR** | *A. constancei* Wendelbo | /Douglasia | USA | Alaska | Alaska (herbarium specimen ALA) | 21/06/2001 | na | 64,96646667 | -143,08573333 | na | 597998 | 597915 |
| **CYH** | *A. cylindrica* DC. | /Argentea | Spain | Pyrenees | Acherito (living plant collection of the Lautaret Alpine Garden) | 29/07/2015 | na | 42,8797458 | -0,7009331 | S. Lavergne, M. Boleda | *3473887* | *3473452* |

| | | | | | | | | | | | | |
|---|---|---|---|---|---|---|---|---|---|---|---|---|
| **AP103** | *A. delphinensis* sp. nov. | /Helvetica | France | Ecrins | Les Rouies, Rébuffat spur | 24/07/2013 | 3330 | 44,862931 | 6,261304 | S. Lavergne, T. Bulle | 5383265 | 5382513 |
| **AP179** | *A. delphinensis* sp. nov. | /Helvetica | France | Ecrins | Pic du Clapier de Peyron | 07/08/2015 | 2995 | 44,9359063 | 6,0796811 | S. Lavergne, J. Charron | 3583357 | 3582818 |
| **AP189** | *A. delphinensis* sp. nov. | /Helvetica | France | Belledonne | Aiguilles Occidentales de l'Argentière | 06/07/2016 | 2900 | 45,2447219 | 6,1316976 | S. Lavergne, M. Boleda | 5647188 | 5646362 |
| **AP25** | *A. delphinensis* sp. nov. | /Helvetica | France | Ecrins | Rateau, S face | 16/07/2009 | 3500 | 44,9982653 | 6,2698452 | S. Ibanez, P. Saccone | 4210072 | 4209441 |
| **PU13** | *A. delphinensis* sp. nov. | /Helvetica | France | Ecrins | Pointe Brevoort, S ridge | 20/08/2014 | 3620 | 44,9669598 | 6,3307025 | S. Lavergne, C. Dentant | 854826 | 854720 |
| **PU3** | *A. delphinensis* sp. nov. | /Helvetica | France | Grandes Rousses | Pic Blanc du Galibier, SE ridge | 07/08/2018 | 2821 | 45,060945 | 6,396312 | S. Lavergne | 2389383 | 2389062 |
| **PU7** | *A. delphinensis* sp. nov. | /Helvetica | France | Ecrins | Sirac, E summit | 26/06/2018 | 3190 | 44,79008 | 6,31754 | C. Dentant | 1971060 | 1970793 |
| **PU8** | *A. delphinensis* sp. nov. | /Helvetica | France | Ecrins | Flambeau, summit | 26/07/2018 | 3550 | 44,92213 | 6,33978 | C. Dentant | 1059968 | 1059824 |
| **PU9** | *A. delphinensis* sp. nov. | /Helvetica | France | Ecrins | Glacier Noir | 26/10/2017 | 2460 | 44,916365 | 6,379336 | C. Dentant | 839898 | 839793 |
| **A_HAL** | *A. halleri* L. | /Halleri | France | Massif Central | Puy Ferrand | 10/06/2016 | na | 45,52394444 | 2,8258333 3 | P. Koutecký | 1208593 | 1208454 |
| **A_HAL_NUR** | *A. halleri* L. | /Halleri | Spain | Pyrenees | Canigou, Plas de Cady | 14/07/2015 | 2500 | 42,5083333 | 2,4333333 | G. Schneeweiss, P. Schönswetter | 2143483 | 2143178 |
| **A_HAU** | *A. hausmannii* Leyb. | /Vitaliana | Italy | Dolomites | Lago Zuoi, Col dei Bos | 04/08/2010 | na | 46,545698 | 12,064176 | A. Moehl | 5486387 | 5485627 |
| **A_HED** | *A. hedraeantha* Griseb. | na | Serbia | Balkans | Mount Midjor, 200m SE from summit | 09/07/2016 | 2150 | 43,3944567 | 22,680548 3 | J. Smyčka, M. Smyčková, D. Požárová | 2118407 | 2118138 |
| **AH_38** | *A. helvetica* (L.) All. | /Helvetica | France | Devoluy | Plateau de Bure | 03/2016 | 2570 | 44,634578 | 5,908646 | E. Hustache, L. Garraud | *1545428* | *1545211* |
| **HE1** | *A. helvetica* (L.) All. | /Helvetica | France | Cerces | Grand Galibier, W ridge | 01/07/2013 | 2690 | 45,0636671 | 6,4124874 | S. Lavergne, F. Boucher | 2182158 | 2181913 |
| **HE2** | *A. helvetica* (L.) All. | /Helvetica | France | Beaufortai n | Col de la Nova | 30/06/2017 | 2770 | 45,653549 | 6,683854 | S. Lavergne, P. Dufour | 947493 | 947361 |
| **HE3** | *A. helvetica* (L.) All. | /Helvetica | France | Aiguilles Rouges | Crochues, bottom of the S ridge | 20/06/2016 | 2770 | 45,9803491 | 6,8729869 | S. Lavergne,S. Ibanez | 945015 | 944878 |
| **HE4** | *A. helvetica* (L.) All. | /Helvetica | France | Cerces | Aretes de la Bruyere | 20/06/2014 | 2450 | 45,0384607 | 6,4765007 | S. Lavergne, C. Dentant | 1006846 | 1006724 |
| **HE5** | *A. helvetica* (L.) All. | /Helvetica | France | Vanoise | La Sechette, limestone bank below the Col des Schistes | 03/08/2016 | 2540 | 45,403219 | 6,7860415 | S. Lavergne, L. Boulangeat | 1725923 | 1725684 |
| **HE6** | *A. helvetica* (L.) All. | /Helvetica | France | Chablais | Dent d'Oche, Plan Champ | 08/06/2018 | 1960 | 46,3498333 | 6,7386783 | S. Lavergne, NB | 2260184 | 2259900 |
| **HE7** | *A. helvetica* (L.) All. | /Helvetica | France | Devoluy | Cabanons de Montmaur | 14/06/2018 | 2450 | 44,6273014 | 5,9167259 | L. Boulangeat, N. Bartalucci | 1149672 | 1149527 |
| **HE8** | *A. helvetica* (L.) All. | /Helvetica | France | Devoluy | Col de Salenton | 07/2018 | 2526 | 46,007879 | 6,854527 | L. Boulangeat | 524902 | 524835 |
| **HE9** | *A. helvetica* (L.) All. | /Helvetica | Switzerland | Bernese Alps | Bella Lui | 31/07/2018 | 2478 | 46,35118333 | 7,4909666 7 | S. Lavergne, F. Boucher, L. Boulangeat | 1354861 | 1354641 |
| **HP2** | *A. helvetica* (L.) All. | /Helvetica | Spain | Pyrenees | Collado de Basibe | 14/07/2014 | 2730 | 42,5409708 | 0,598154 | S. Lavergne | *2188943* | *2188665* |
| **D_IDA** | *A. idahoensis* (Douglass M.Hend.) Cubey | /Douglasia | USA | Idaho | Boise National Forest (herbarium specimen ID) | 07/07/2006 | na | 44,305361 | - 115,74180 6 | M. Mancuso | 767656 | 767546 |
| **A_KOM** | *A. komovensis* Schönsw. & Schneew. | /Vitaliana | Kosovo | Balkans | Mount Marjashi | 15/08/2010 | na | 42,6538889 | 20,111666 7 | G. Schneeweiss | 2119153 | 2118878 |
| **A_LAC** | *A. lactea* L. | /Argentea | Switzerland | Bernese Alps | Gantrisch | 08/08/2009 | na | 46,711958515 4824 | 7,4403288 683004 | A. Moehl | 1603477 | 1603282 |
| **A_LAG** | *A. laggeri* A.Huet. | /Halleri | Spain | Pyrenees | Portarro d'Espot, | 26/07/2016 | na | 42,57908333 | 0,9743333 | S. Lavergne, C. Roquet, | 2043282 | 2043047 |

| | | | | | | | | | | | | |
|---|---|---|---|---|---|---|---|---|---|---|---|---|
| | | | | | Encantats | | | | 3 | J. Smycka | | |
| **A_MAT** | *A. mathildae* Levier | /Argentea | Italy | Appenins | Gran Sasso, 200m below summit | 15/08/2016 | 2700 | 42,4717931 | 13,5621067 | J. Smyčka, J.Pilátová, M. Kolářová | 696691 | 696593 |
| **A_OBT** | *A. obtusifolia* All. | /Argentea | France | Grandes Rousses | Château-Noir, Clavans-en-Haut-Oisans | 14/07/2009 | na | 45,103806 | 6,133339 | J.-C. Villaret | 1811959 | 1811724 |
| **OBT** | *A. obtusifolia* All. | /Argentea | Switzerland | Valais | Schwarsee, along the trail to the Matterhorn | 01/08/2018 | 2638 | 45,9893 | 7,70455 | S. Lavergne, F. Boucher, L. Boulangeat | 1131611 | 1131449 |
| **AP_231** | *A. pubescens* DC. | /Helvetica | France | Chablais | La Pointe, col de Ratti | 08/06/2018 | 1890 | 46,1873773 | 6,6212335 | S. Lavergne, N. Bartalucci | 1429891 | 1429711 |
| **AP_238** | *A. pubescens* DC. | /Helvetica | France | Chablais | Dent d'Oche, Plan Champ | 08/06/2018 | 1960 | 46,3498333 | 6,7386783 | S. Lavergne, N. Bartalucci | 2779144 | 2778777 |
| **PU1** | *A. pubescens* DC. | /Helvetica | France | Devoluy | Cabanons de Montmaur | 14/06/2018 | 2450 | 44,6273014 | 5,9167259 | L. Boulangeat, N. Bartalucci | 524900 | 524833 |
| **PU5** | *A. pubescens* DC. | /Helvetica | Switzerland | BerneseAlps | Bella Lui | 31/07/2018 | 2486 | 46,35203333 | 7,49098333 | S. Lavergne, F. Boucher, L. Boulangeat | 3031633 | 3031251 |
| **PY2** | *A. pyrenaica* Lam. | /Halleri | Spain | Pyrenees | Puertolas | 06/09/2010 | na | 42,58513 | 0,07205 | S. Lavergne | 582387 | 582313 |
| **PY3** | *A. pyrenaica* Lam. | /Halleri | Spain | Pyrenees | Cabana del Foradet, Posets | 23/07/2003 | na | 42,589783 | 0,411455 | G. Schneeweiss | 4150413 | 4149876 |
| **A_RIO** | *A. rioxana* A.Segura | /Halleri | Spain | Sierra Iberica | Pico San Lorenzo, La Rioja | 02/07/1905 | 2200 | 42,2333333 | -2,95 | G. Schneeweiss | 2994310 | 2993943 |
| **AA32** | *A. saussurei sp. nov.* | /Helvetica | France | Mont Blanc | Tour Ronde | 06/2017 | 3609 | 45,842 | 6,91 | S. Ibanez | 3346547 | 3346072 |
| **AP217** | *A. saussurei sp. nov.* | /Helvetica | France | Vanoise | Grande Gliere, bottom of SE ridge | 04/07/2017 | 3150 | 45,41113 | 6,793562 | S. Lavergne, M. Boleda | 2128887 | 2128552 |
| **AX_19** | *A. saussurei sp. nov.* | /Helvetica | France | Mont Blanc | Ridge of the Grandes Autannes | 21/06/2016 | 2520 | 46,020779 | 6,9787327 | S. Lavergne, C. Dentant, S. Ibanez | 3054398 | 3053975 |
| **AX_47** | *A. saussurei sp. nov.* | /Helvetica | France | Mont Blanc | Dent du Géant | 24/06/2016 | 3605 | 45,8605028 | 6,9486459 | S. Lavergne, C. Dentant, S. Ibanez | 2488564 | 2488236 |
| **AX_84** | *A. saussurei sp. nov.* | /Helvetica | France | Thabor | Pic du Thabor | 09/2017 | 3100 | 45,1194361 | 6,5618778 | S. Ibanez | 3003280 | 3002896 |
| **AX17** | *A. saussurei sp. nov.* | /Helvetica | France | Mont Blanc | Ridge of the Grandes Autannes | 21/06/2016 | 2520 | 46,020779 | 6,9787327 | S. Lavergne, C. Dentant, S. Ibanez | 4991080 | 4990335 |
| **IN2** | *A. saussurei sp. nov.* | /Helvetica | Italy | Gran Paradiso | Herbetet | 20/08/2018 | 3600 | 45,54343 | 7,26932 | N. Bartalucci | 2197512 | 2197210 |
| **IN5** | *A. saussurei sp. nov.* | /Helvetica | France | Mont Blanc | Aiguille du Tour | 06/2013 | 3400 | 45,994286615 | 7,010692958 | S. Ibanez | 2321962 | 2321667 |
| **IN6** | *A. saussurei sp. nov.* | /Helvetica | France | Mont Blanc | Grands Mulets | 04/2017 | 2997 | 45,866 | 6,861 | S. Ibanez | 617611 | 617523 |
| **AP181** | *A. saussurei x pubescens* | /Helvetica | France | Aiguilles Rouges | Crochues, bottom of the S ridge | 20/06/2016 | 2770 | 45,9803491 | 6,8729869 | S. Lavergne,S. Ibanez | 3164519 | 3164043 |
| **IN1** | *A. saussurei x pubescens* | /Helvetica | Switzerland | Valais | Grand Combin, W ridge of Valsorey | 23/09/2018 | 3950 | 45,937593 | 7,299102 | S. Lavergne | 3805753 | 3805297 |
| **IN3** | *A. saussurei x pubescens* | /Helvetica | Italy | Mont Blanc | Freney Pilar | 09/2014 | 4050 | 45,825249085 | 6,874417227 | S. Ibanez | 511122 | 511067 |
| **IN4** | *A. saussurei x pubescens* | /Helvetica | France | Mont Blanc | Goûter | 06/2013 | 3400 | 45,853713511 | 6,826310074 | S. Ibanez | 1165710 | 1165565 |
| **A_SEP** | *A. septentrionalis* L. | outroup | Switzerland | Grison Alps | Zernez | 17/06/2009 | na | 46,6987914250095 | 10,0912459230913 | A. Moehl | 3238709 | 3238303 |
| **AP_226** | *A. vesulensis* sp. nov. | /Helvetica | Italy | Cottian Alps | Monte Viso | 18/07/2017 | 3750 | 44,66756 | 7,08978 | S. Lavergne, J. Smycka | 2512943 | 2512636 |
| **AP223** | *A. vesulensis* sp. nov. | /Helvetica | Italy | Cottian Alps | Monte Viso | 18/07/2017 | 3750 | 44,66756 | 7,08978 | S. Lavergne, J. Smycka | 2418794 | 2418420 |
| **PU10** | *A. vesulensis* sp. nov. | /Helvetica | Italy | Cottian Alps | Monte Viso | 07/2013 | 3450 | 44,66364579 | 7,092131193 | S. Ibanez | 1433757 | 1433571 |
| **PU11** | *A. vesulensis* sp. nov. | /Helvetica | Italy | Cottian | Monte Viso | 18/07/2017 | 3750 | 44,66756 | 7,08978 | S. Lavergne, J. Smycka | 2098451 | 2098181 |

| | | | | | | | | | | | | |
|---|---|---|---|---|---|---|---|---|---|---|---|---|
| | | | | Alps | | | | | | | | |
| **PU12** | *A. vesulensis* sp. nov. | /Helvetica | Italy | Cottian Alps | Monte Viso | 18/07/2017 | 3750 | 44,66756 | 7,08978 | S. Lavergne, J. Smycka | 1410153 | 1409988 |
| **A_VIT_CIN** | *A. vitaliana* (L.) Lapeyr. | /Vitaliana | France | Ecrins | Galibier | 07/03/2009 | na | 45,0640078097227 | 6,40852715398399 | R. Douzet, C. Roquet, M. Boleda | 3574580 | 3574107 |
| **A_VIT_SES** | *A. vitaliana* (L.) Lapeyr. | /Vitaliana | Italy | Dolomites | Above Rifugio Bindel | 21/06/2012 | na | 46,47697 | 11,841838 | A. Moehl | 5421908 | 5421150 |
| **A_VIT_VIT** | *A. vitaliana* (L.) Lapeyr. | /Vitaliana | France | Devoluy | Rocher Rond | 16/07/2010 | na | 44,7067 | 5,80042 | J.-C. Villaret | 3309432 | 3308984 |
| **A_WUL** | *A. wulfeniana* Hand.-Mazz. | /Vitaliana | Austria | Eastern Alps | Triebental | 15/07/2019 | na | 47,3648 | 14,54401 | S. Latzin | 2172248 | 2171963 |
| **LYS_NUM** | *Lysimachia nummularia* L. | outroup | Switzerland | Berne canton | Suberg | 25/05/2009 | na | 47,0637413 | 7,33318933 | S. Eggenberg | 720764 | 720661 |
| **PHV** | *Primula hirsuta* All. | outroup | France | Belledonne | Col de Villonet | 14/07/2018 | 2460 | 45,3179 | 6,1946 | F. Boucher | 705426 | 705338 |

**Table S1.** List of all samples used in this study with collection and sequencing information.

**SUPPLEMENTARY INFORMATION FOR :**

### Discovery of cryptic plant diversity in one of the harshest environments: the rooftops of the Alps

by

Boucher, F.C., Dentant, C., Ibanez, S., Capblancq, T., Boleda, M., Boulangeat, L., Smyčka J., Roquet C., Noûs C. & Lavergne, S

#### 1. Genomic data acquisition

*1.1 ddRAD-seq protocol*

We used a modified version of the protocol described in [1]. A total of 150 ng of DNA template from each individual were double-digested with 10 units each of PstI and MspI (New England Biolabs Inc.) at 37°C during one hour in a final volume of 34 µl using the CutSmart buffer provided with the enzymes. Digestion was further continued together with ligation of P1 and P2 adapters (see Peterson et al. 2012) by adding 10 units of T4 DNA ligase (New England Biolabs Inc.), adapters P1 and P2 in 10-fold excess (compared to the estimated number of restriction fragments) and 1 µl of 10 mM ribo-ATP (New England Biolabs Inc.) in each sample. This simultaneous digestion-ligation reaction was performed on a thermocycler using a succession of 60 cycles of 2 min at 37°C for digestion and 4 min at 16°C for ligation. An equal volume mixture of all the digested-ligated fragments were purified with Agencourt AMPure XP beads (Beckman Coulter, France). Fragments were size-selected between 200 and 600 bp on a Pippin Prep. The ddRAD libraries obtained were amplified using the following PCR mix: a volume of 20µl with 2 µl of DNA template, 10 mM of dNTPs, 2 µM of each PCR probe (Peterson et al. 2012) and 2 U/µl of Taq Phusion-HF (New England Biolabs Inc.); and the following PCR cycles: an initial denaturation at 98°C for 30 sec; 15 cycles of 98°C for 10 sec, 66°C for 30 sec and 72°C for 1 min; followed by a final extension period at 72°C for 10 min. The four amplified ddRAD libraries were purified with a QIAgen MinElute PCR Purification Kit (Qiagen, Germany) and then sequenced on half a lane of Illumina Hi-Seq 2500 2x125 (Fasteris SA, Switzerland).

*1.2 Bioinformatic assembly of ddRAD tags*

All sequenced ddRAD-seq tags (~275 million reads, see details in Table S1) were assembled using the program ipyrad (https://github.com/dereneaton/ipyrad) using the following parameters :

- a maximum of 5 bases with quality <20 per read

- a phred score offset of 33 ; a minimum depth of 6 reads for both statistical and majority-rule base calling

- a maximum cluster depth of 10,000 within individual samples

- a clustering threshold of 0.88 when using *de novo* assembly (only used for sensitivity analyses in our case, see below)

- a minimum length of 35bp after quality trimming of reads

- a maximum of 2 alleles per site in consensus sequences

- a maximum of 5 uncalled bases per *locus*

- a maximum of 8 heterozygous sites per *locus*

- a maximum of 20 SNPs per *locus* (which was never reached)

- a maximum of 8 indels per *locus*

Rather than relying on *de novo* assembly of these *loci*, we preferred to align them on the reference genome of another species from the Primulaceae family: *Primula veris* L. [2]. This reference genome has already been used successfully in a phylogenomic study of a group of plants that had diverged > 20 Ma from it [3] and in our case this reference genome has been estimated to have diverged c. 33 Ma from *Androsace* sect. *Aretia* [4]. While we lost a large number of loci by aligning to this rather distant reference genome compared to *de novo* assembly (2,700 vs. > 50,000 loci recovered), we preferred to rely on this reduced dataset in order to avoid paralogs as much as possible. We however verified that a dataset containing both loci aligned to the reference genome and *de novo* assembly of loci that did not align to the reference genome (‘denovo+reference’ option in ipyrad) provided qualitatively similar results (see below). Before combining them for further analyses, we checked that technical replicates of our various ddRAD-seq libraries and sequencing runs produced comparable data. To do so we inferred a phylogeny of the 31 individuals for which we had replicates (97 replicates in total, up to four for a single individual) using maximum-likelihood (hereafter, ‘ML’) with concatenation as detailed below. The resulting phylogram showed that all replicates from the same individual formed clades with internal branch lengths much shorter than branches connecting different individuals (Fig. S1), and we thus combined all sequence data from different replicates of the same individual for subsequent analyses.

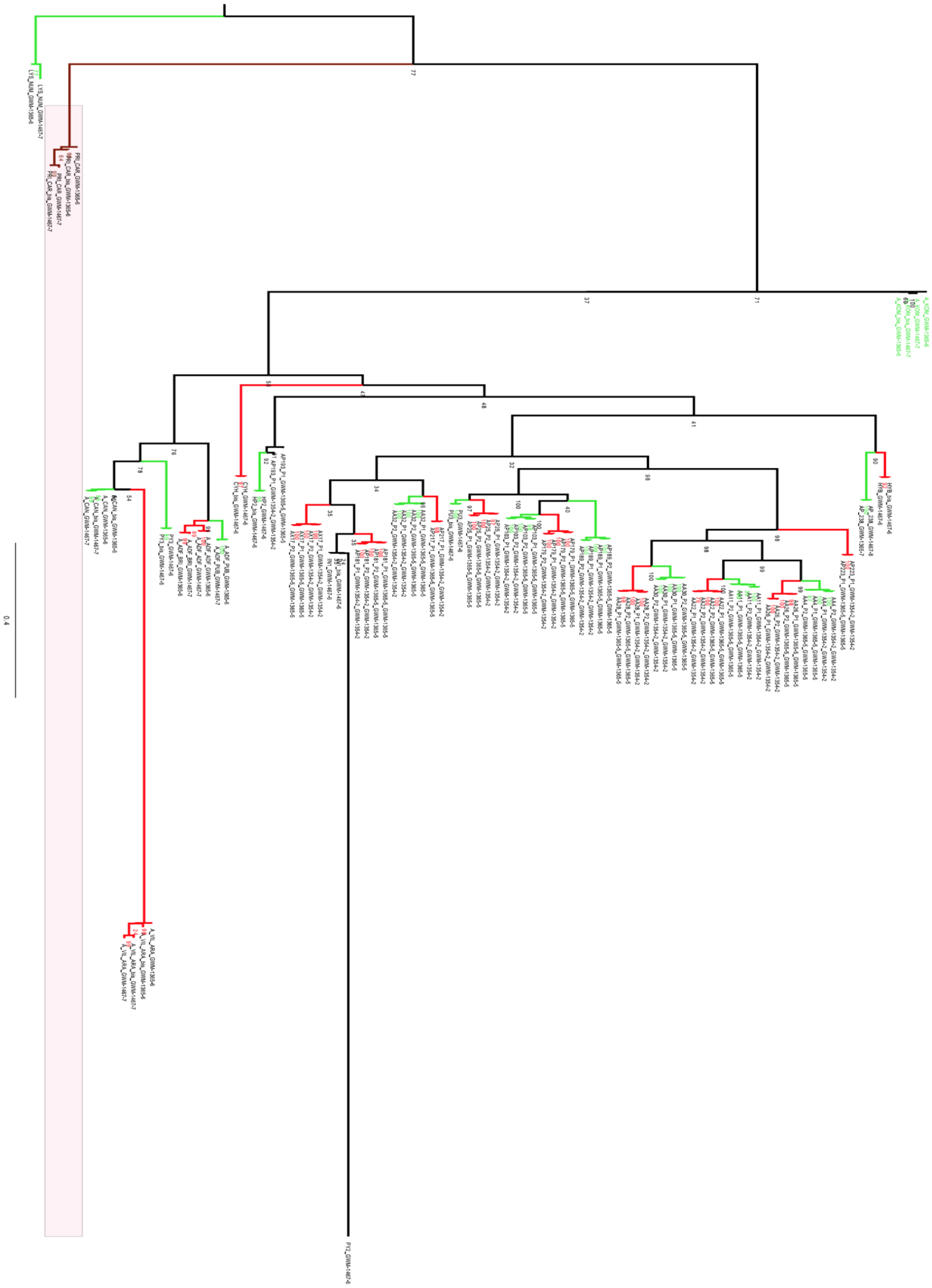

**Fig. S1. Repeatability of different libraries and sequencing runs.** The figure shows the ML phylogeny of 97 replicates, obtained using RAxML [5] under de GTR+GAMMA model with 100 bootstrap replicates. Clusters of technical replicates from the same individual are shown in the same color and boostrap is shown at nodes.

#### 2. Phylogenetic inference

##### *2.1. Phylogenetic relationships*

Phylogenetic relationships within *Androsace* sect. *Aretia* were inferred using two different approaches. ML inference was first carried out using IQ-TREE [6] with all ddRADseq loci concatenated (314,363 bp). The best model of DNA substitution for this dataset, TPM2u+F+R3, was selected among 286 possible ones according to BIC, and 1000 ultrafast bootstrap replicates [6] were used to measure node support on this phylogeny. In addition to this approach, we also inferred a species tree under the multi-species coalescent in order to account for incomplete lineage sorting and incongruence between gene trees [7]. This was done by selecting unlinked SNPs from the initial matrix, *i.e.* SNPs that were either located on different contigs of the reference genome or SNPs located on the same contig but >10,000 bp apart. These 2,461 SNPs were then used in the program SVDquartets [8], assembling all possible quartets and using 100 bootstrap replicates to measure branch support. In both cases, trees were rooted thanks to the inclusion of three outgroups from the Primulaceae family, located at increasing phylogenetic distances from our ingroup: *A. septentrionalis* L., *Primula hirsuta* All., and *Lysimachia nummularia* L..

Both phylogenetic inference methods were largely congruent and led to a highly supported phylogenetic hypothesis (Fig. S2, Fig. S3). They first recovered the split of sect. *Aretia* into two large clades [9]: one of them is made up of species from North-America as well as various mid-elevation mountain ranges of Western Europe (clade /Dicranothrix), and the second one mostly consists of high-altitude cushion-forming species from the Alps and adjacent mountain ranges (clade /Eu-Aretia). Furthermore, ML and species tree inference supported the existence of five subclades. One of them, /Douglasia, corresponds to the former genus *Douglasia* Lindley and the four others are named after their most widespread species: /Halleri, /Vitaliana, /Argentea, and /Helvetica. Among these clades, /Douglasia received 100% bootstrap support in both analyses, while /Vitaliana and /Helvetica were also strongly supported as monophyletic (bootstrap support >

83% in both analyses). Of the twelve species for which multiple individuals were included, only *A. alpina* and *A. halleri* were not recovered as being monophyletic (Fig. S2). In the first case, the apparent paraphyly of *A. alpina* is weakly supported, and subsequent analyses including more samples and focused on clade /Helvetica clearly support the monophyly of this species (see below). In the second case, the paraphyly of *A. halleri* confirms previous phylogenetic results suggesting that *A. rioxana* is nested within it [10].

These results represent a major improvement over previous studies of *Androsace* sect. *Aretia*. While this group, and more broadly the genus *Androsace*, have been the focus of several phylogenetic studies, all of them have relied on traditional Sanger sequencing with emphasis on chloroplastic markers and present conflicts between DNA and morphology [9,11]. Our use of ddRADseq data solves all of these issues and recovers five clades which composition aligns well with both geography and morphology. This comes from the vast amount of DNA sequence data analyzed here (> 10 million bp for the largest dataset), from the fact that it is mostly nuclear DNA, but also from the use of the multispecies coalescent allowed by this kind of data that strengthens our confidence in phylogenetic inferences. Such an improved knowledge of phylogenetic relationships using multilocus sequence data had already been obtained in a closely related group of plants from the European Alpine System, *Primula* sect. *Auricula* [3].

**Fig. S2.**

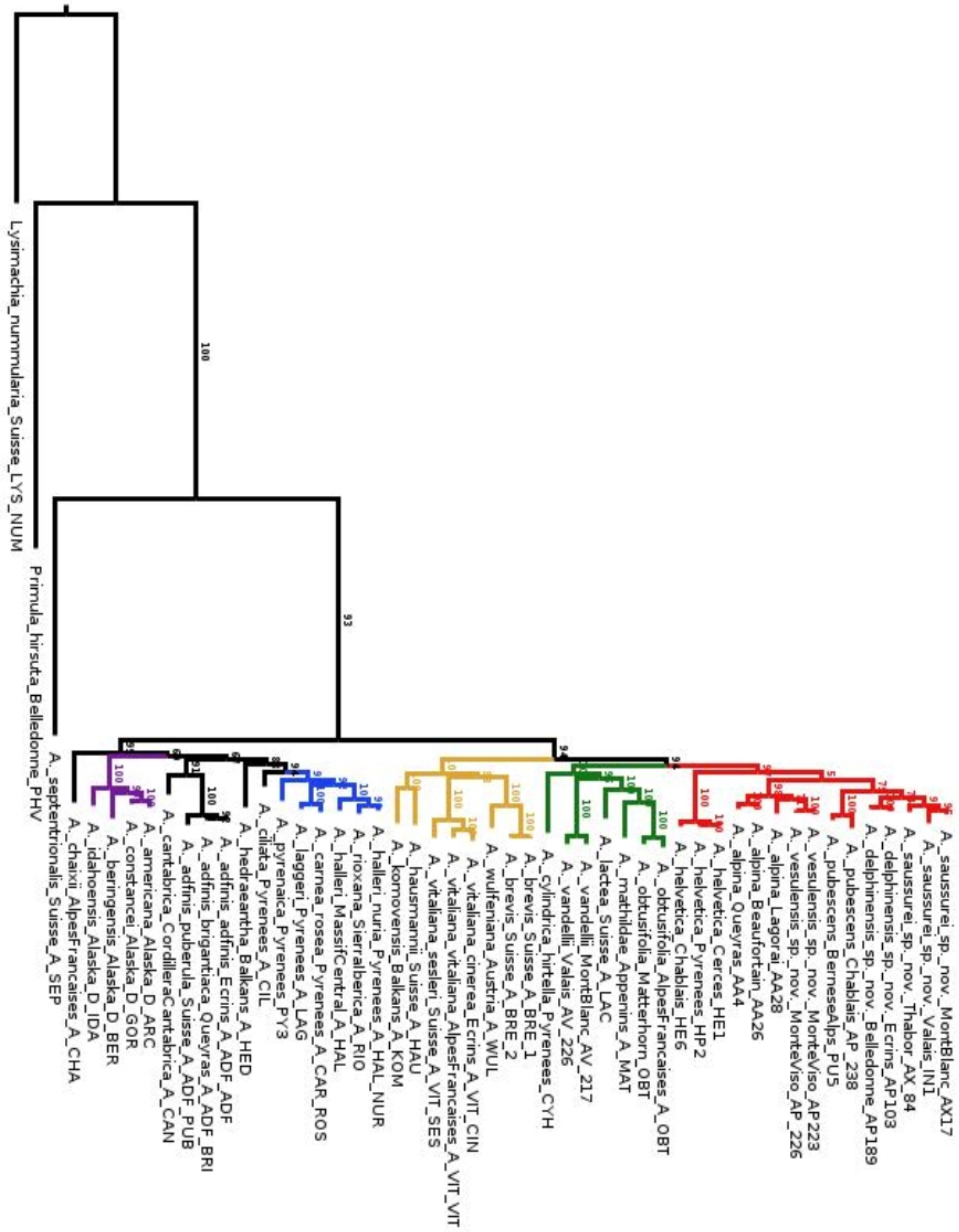


**Phylogenetic relationships within *Androsace* sect. *Aretia*** inferred from ML using a concatenation of all ddRAD tags mapped to the reference genome (314,363 bp). Bootstrap support is shown above branches and the five clades that are supported by both this ML inference and a species tree analysis have been colored and labeled.

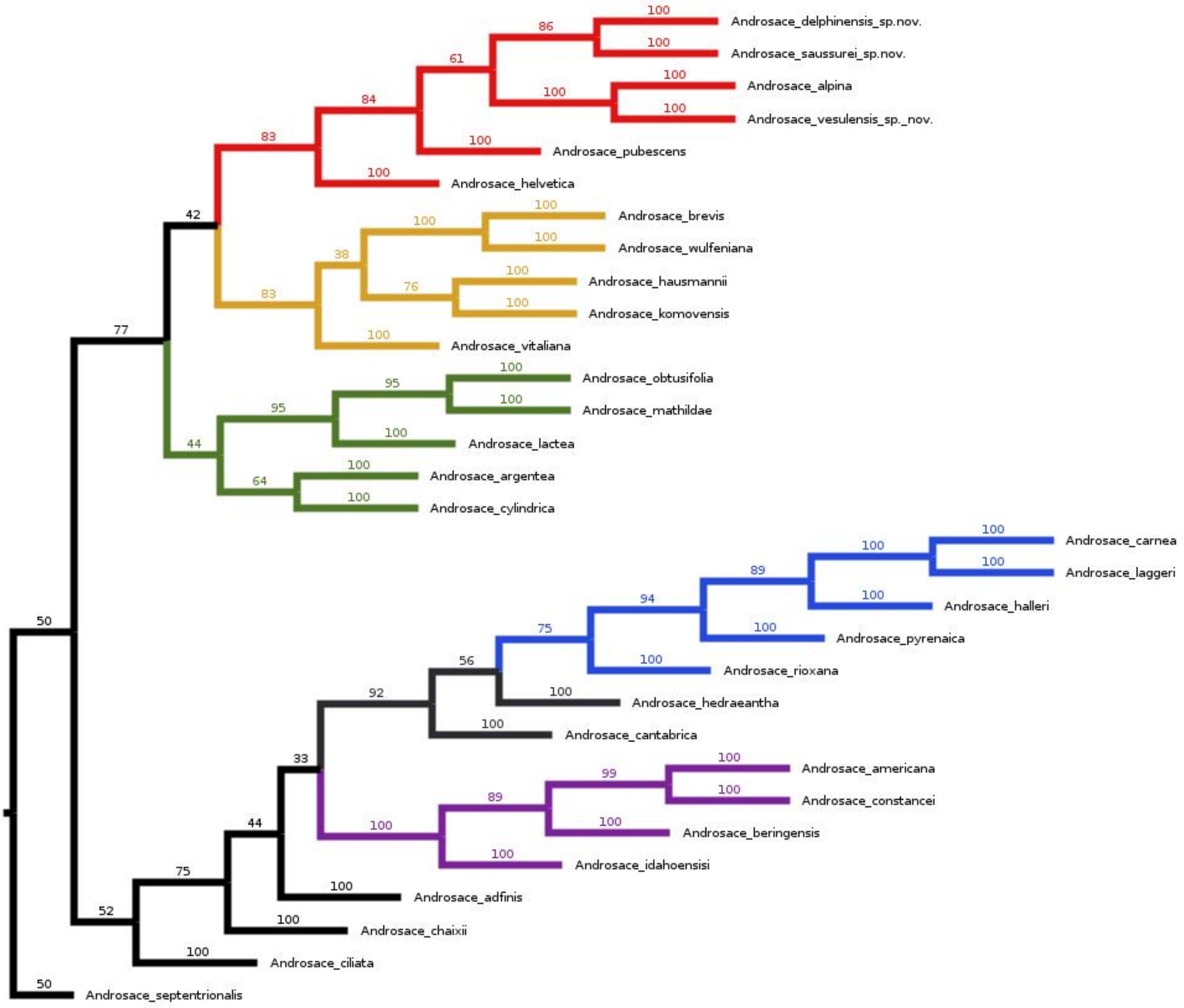


**Fig. S3. Phylogenetic relationships within *Androsace* sect. *Aretia*** inferred using species tree inference from 2,461 unlinked SNPs mapped to the reference genome. Bootstrap support is shown above branches and the five clades that are supported by both this species tree analysis and ML inference have been colored and labeled.mapped to the reference genome (314,363 bp)

*2.2. Divergence time estimation*

In order to estimate the time frame of diversification of *Androsace* sect. *Aretia*, we then proceeded to date the phylogram obtained thanks to ML using penalized likelihood [12]. To do so, we compared seven different values of the smoothing parameter λ (0.001, 0.01,0.1,0.5,1,2,10), and retained the single one that gave the lowest ΦIC [13]. We calibrated two nodes of the phylogeny using median ages estimates from a previous study of Primulaceae [4]: the crown node of *Androsace* sect. *Aretia* (8.16 Ma), as well as the divergence between the genera *Androsace* and *Primula* (33.3 Ma). While secondary calibration is generally prone to error propagation [14], this was the only analytically-tractable solution here given the size of the phylogenomic dataset, which precluded more computationally intensive methods.

Results suggested that all five subclades might have originated before the Pleistocene, most likely in the Pliocene (Fig. S4). The youngest one of them is /Halleri (2.33 Ma) and the oldest is /Vitaliana (5.60 Ma). High-alpine cushion species of the /Helvetica clade were estimated to have originated at 4.66 Ma.

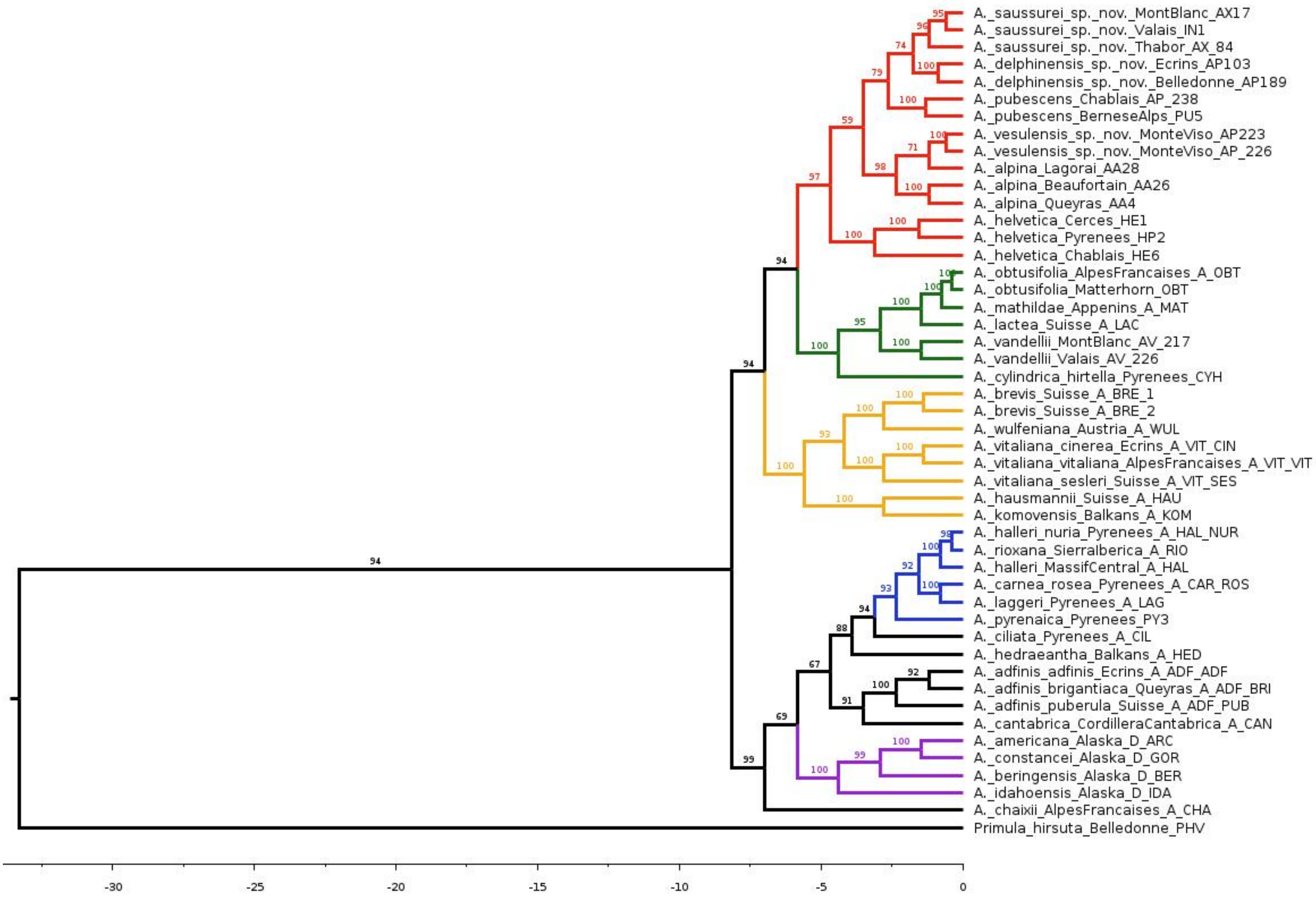


**Fig. S4. Time-calibrated phylogeny of *Androsace* sect. *Aretia*** based on a concatenation of 2,700 ddRADseq tags mapped to the reference genome (314,363 bp) and dated using penalized likelihood. Time is shown in million years on the x-axis and bootstrap support is shown above branches. The five clades that are supported by both this ML inference and a species tree analysis have been colored.

#### 3. Systematics of the /Helvetica clade

In order to revise the systematics of /Helvetica and to test the putative species status of newly discovered taxa, we used an integrative taxonomic approach combining genomics, morphology, geography, and bedrock affinities.

##### *3.1. Assembly of ddRAD tags and SNP filtering*

We started by doing a different SNP calling for the 51 individuals of the /Helvetica clade plus one outgroup individual from A. vitaliana, using the same pipeline as described above for the whole section *Aretia*, and in particular the same parameters in ipyrad. This resulted in 23,780 loci aligned to the reference genome, containing a total of 276,745 bp and 7,806 SNPs.

In order to use population genetic measures on this rather recently diverged clade (4.66 Ma following the estimation done above), we strictly filtered this initial SNP dataset: we only kept SNPs that had less than 40% missing data, that had minor allele frequencies >4% (i.e., alleles genotyped at least three times, thus in two individuals minimum), and that were unlinked (*i.e.*, located either on different contigs of the reference genome or > 10,000 bp apart on the same contig). This yielded a final dataset of 381 SNPs. Although this number can be considered low compared to other studies that tackled the same questions (e.g., 15, 16), we preferred working with these high-quality SNPs only rather than using many more SNPs. SNPs datasets obtained from de novo assembly of ddRADseq tags may indeed be prone to include PCR or sequencing errors as well as paralogs. However, for analyses that could handle huge datasets (*i.e.*, ML phylogenetic inference and genetic clustering), we did confirm results obtained from this reference mapping dataset with a ‘denovo+reference’ dataset as described above, which contained a total of 10,552,054 bp, and 2,234 SNPs when filtering with the same parameters as above (except that SNPs are now considered unlinked if they are present on different de novo-assembled RAD tags). Results from both analyses were highly similar to those obtained using the reference mapping dataset (see below).

*3.2. Genetic structure*

Genetic clusters were then inferred with no *a priori* using an efficient algorithm aimed at estimating individual ancestry: sNMF [17]. Given that /Helvetica currently comprises three recognized species and since we expected up to three new taxa, we ran the program for a number of genetic clusters K ranging from 1 to 10 with 1,000 iterations and 20 repetitions per value of K. After verifying that different values of the regularization parameter α gave similar results we chose to use a value of 50, and the optimal number of clusters was chosen based on the cross-entropy of the best run for each value of K [17].

Using either only the 381 SNPs obtained after strict filtering of reads aligned to the reference genome (Fig. S5) or the 2,234 SNPs obtained after strict filtering of reads from the 'denovo+reference' dataset (Fig. S6) we inferred the same number of K=7 optimal clusters. Ancestry proportions of the different individuals are similar for both SNP datasets, when looking from 5 up to the optimal 7 clusters (Fig. S5, S6), and thus lead to the same delimitation of genetic clusters.

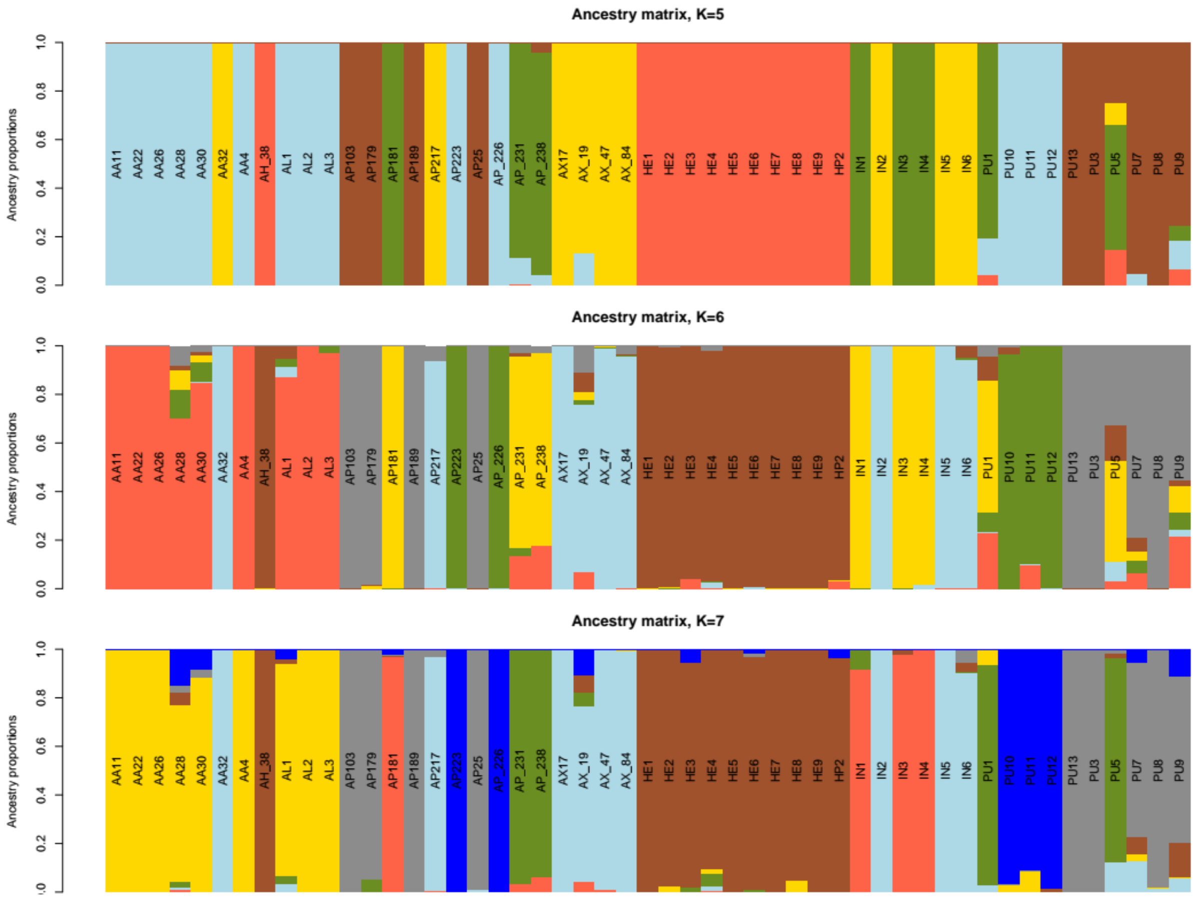


**Fig. S5. Ancestry proportions for the 51 individuals of the /Helvetica clade** , obtained using the most restricted dataset with only SNPs mapped onto the reference genome . Proportions are shown for K=5, K=6, and K=7 clusters (the optimal number here). Individual codes correspond to those given in Table S1, but be careful that the colors used to picture different genetic clusters differ from those in Fig. 2, S6, S7 and S8.

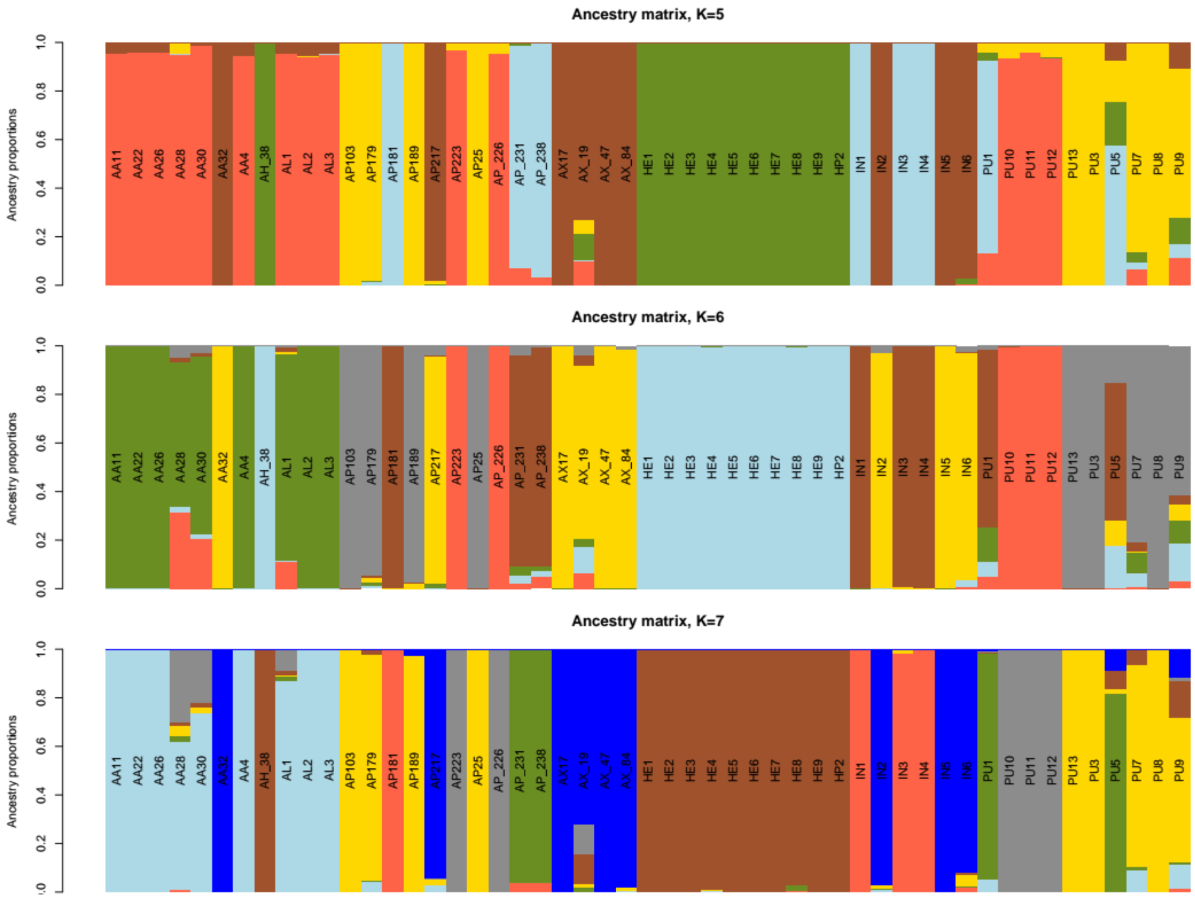


**Fig. S6. Ancestry proportions for the 51 individuals of the /Helvetica clade** , obtained using SNPs from the 'de novo + reference' dataset. Proportions are shown for K=5, K=6, and K=7 clusters (the optimal number here). Individual codes correspond to those given in Table S1 and are shown in the same order as in Fig. S5, but be careful that the colors used to picture different genetic clusters differ from those in Fig. 2, S5, S7 and S8.

Samples assigned to the three currently recognized species in /Helvetica were distributed among the seven inferred clusters as follows:

- all individuals assigned to *A. helvetica,* from both the Alps and the Pyrenees, were assigned to the same cluster (the brown one in Fig 3 & S5).

- individuals initially assigned to *A. alpina* were split into two distinct clusters: one comprising individuals from Monte Viso only (hereafter referred to as *A. vesulensis sp. nov.* and pictured in light blue in Fig. 3 & S5) and another one comprising all other individuals from across the Alps (*A. alpina*, pictured in dark blue in Fig. 3 & S5).

- individuals initially assigned to *A. pubescens* were split into three clusters. One included all individuals growing on limestone from the French and Swiss Alps (hereafter, *A. pubescens,* pictured in green in Fig. 3 & S5). The second cluster included individuals growing on acidic substrates in the Mont Blanc and neighboring ranges (hereafter referred to as *A. saussurei sp. nov.,* pictured in yellow in Fig. 3 & S5). Finally, the third cluster included individuals growing on acidic substrates in the Central French Alps (hereafter referred to as *A. delphinensis sp. nov.,* pictured in grey in Fig. 3 & S5).

Finally, the seventh, and last, genetic cluster corresponded to four individuals from the Mont Blanc and neighboring ranges that had intriguing leaf trichomes not seen in any other individual: these were antler-shaped as typically seen in some species (see *Determination key* below) but were much longer, as seen in the case of simply bifurcated trichomes. We thus tested whether this cluster could result from introgression between *A. saussurei* sp. nov. and *A. pubescens* as measured by Patterson's D-statistic, using the ABBA-BABA test [18]. Based on ML phylogenetic results (see below), this test was run with *A. alpina* as the outgroup, and *A. saussurei* sp. nov. plus *A. pubescens* as ingroup taxa on top of the putatively introgressed cluster. Two different topologies were tested, with the putatively introgressed cluster as P2 in the typical ABBA-BABA terminology, and exchanging the two other ingroup taxa between P1 and P3 positions (Fig. S7). In both cases, 1,000 bootstrap replicates of the D-statistic were obtained by randomly sampling one individual in each of these four lineages and we then calculated a p-value for the test by counting the proportion of these replicates that exceeded 0. The two alternative phylogenetic topologies showed significant signals of introgression (p-value<0.043 for both), supporting an hybrid origin of this group of individuals from the Mont Blanc area, where the two parent taxa (*A. saussurei sp. nov.* and *A. pubescens* both occur).

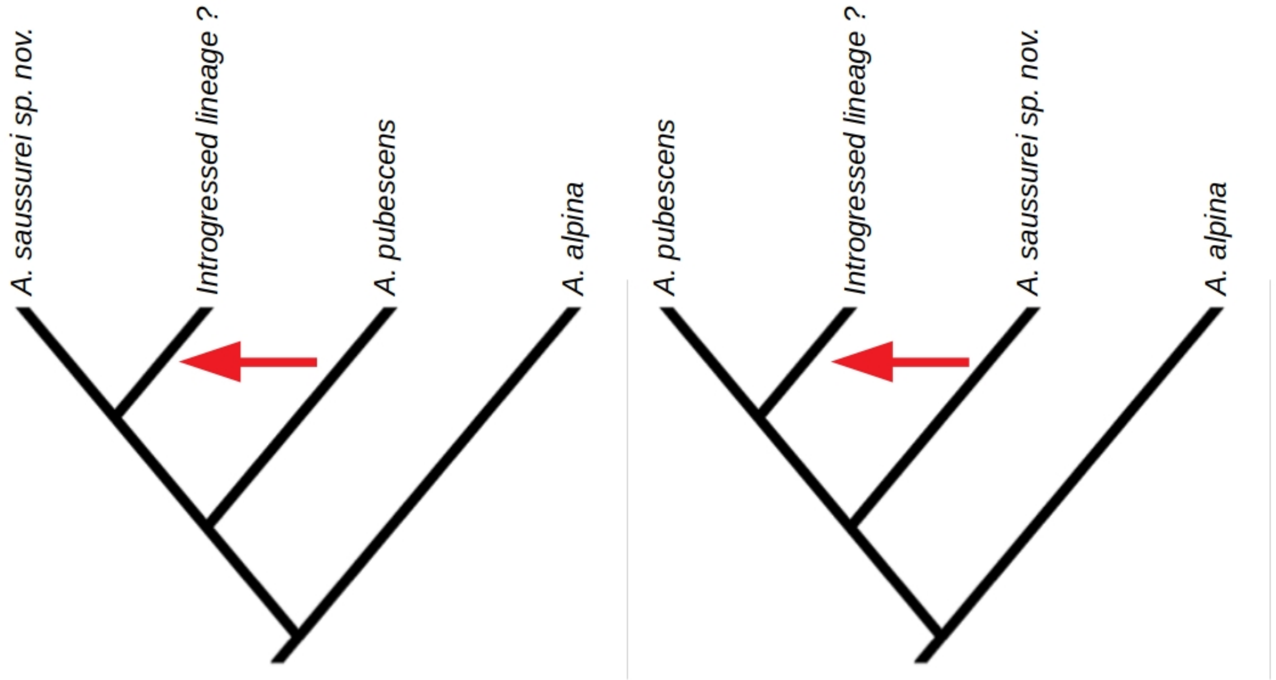


**Fig. S7. Test for introgression in one of the seven genetic clusters.** These graphs show the two phylogenetic topologies that were used to test for introgression in one of the seven clusters that contained four individuals with an intermediate trichome morphology (the ones colored in pink for K=7 in Fig. 3, S5 & S6). In both cases, the introgression event that is tested by the ABBA-BABA test is shown using the red arrow.

We complemented this clustering analysis with the inference of phylogenetic relationships within clade /Helvetica. To do so we inferred an ML phylogeny of all 51 individuals using concatenation of all 276,745bp. The best sequence evolution model selected according to BIC by IQ-TREE [6] was HKY+F+R2 and 1000 ultrafast bootstrap samples were generated to measure clade support in the ML phylogeny. In this phylogeny, the seven distinct genetic clusters corresponded to seven clades that all received 100% bootstrap support, except that the introgressed lineage was nested within *A. saussurei sp. nov.* (Fig. S8).

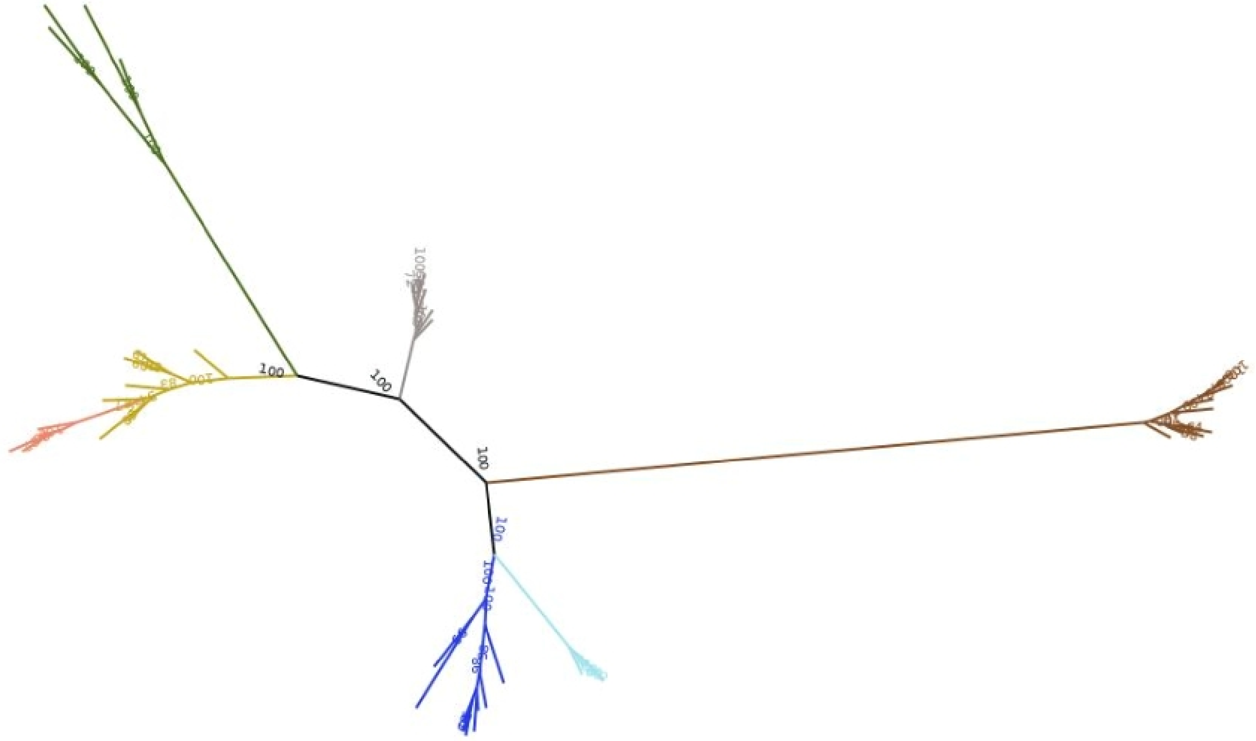

**Fig. S8. Unrooted phylogeny of 51 individuals of the /Helvetica clade** based on a concatenated DNA matrix of 276,745 bp obtained from the alignement of ddRAD tags on the reference genome of *Primula veris,* and inferred using ML. Bootstrap is shown at nodes, but only visible for the deepest ones in the tree. Tip labels have been omitted for clarity and colors of the different clades (or grade in the case of *A. saussurei sp. nov.*, in yellow) match those in Fig. 3, S5 & S6.

Results were qualitatively unchanged when using a much larger (but potentially more error prone) ‘de novo + reference’ dataset. The best sequence evolution model was again HKY+F+R2 and 1000 ultrafast bootstrap samples were generated to measure clade support in the ML phylogeny. The seven distinct genetic clusters again corresponded to seven clades that all received 100% bootstrap support, except that the introgressed lineage was nested within *A. saussurei sp. nov.* (Fig. S8).

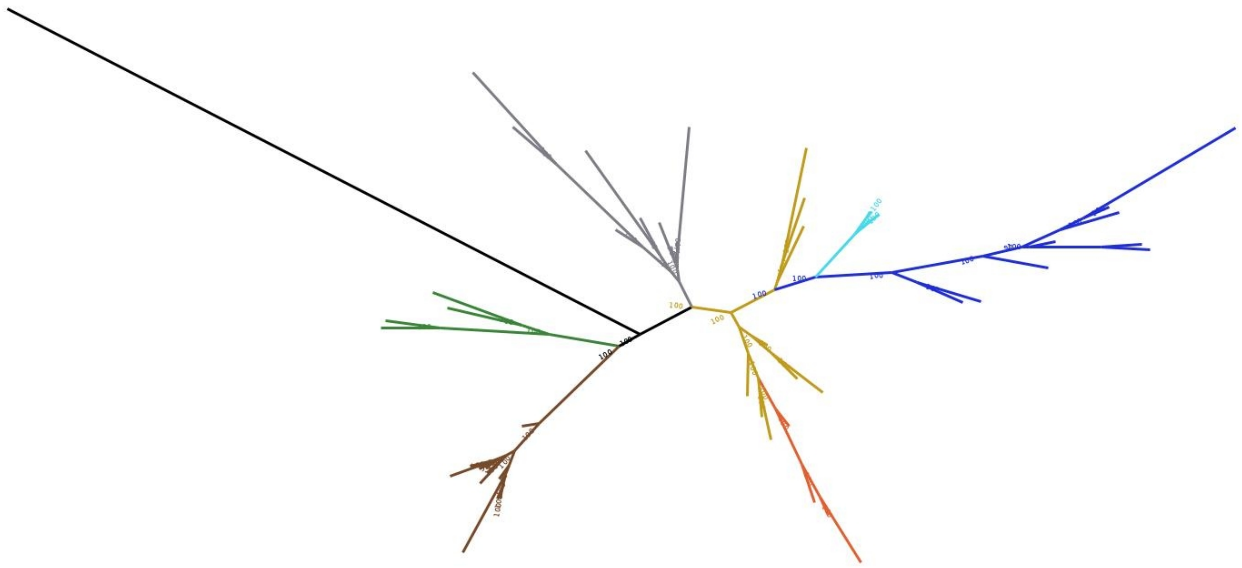

**Fig. S9. Unrooted phylogeny of 51 individuals of the /Helvetica clade** based on a concatenated DNA matrix of 10,552,054 bp obtained from the alignement of ddRAD tags from the ‘de novo + reference’ dataset, and inferred using ML. Bootstrap is shown at nodes, but only visible for the deepest ones in the tree. Tip labels have been omitted for clarity and colors of the different clades (or grade in the case of *A. saussurei sp. nov.*, in yellow) match those in Fig. 3, S5 & S6. An additional individual from *A. vitaliana* (not a member of /Helvetica) is shown in black but the tree is drawn unrooted for comparison with Fig. S8.

*3.3. Species delimitation and taxonomic status*

We tested the species status of the five clusters that did not correspond to already recognized species using the explicit approach of Bayes factor delimitation (BFD*,19) that relies on the SNAPP method to directly estimate a species tree from unlinked biallelic markers without explicitly reconstructing gene trees [20]. To statistically test for alternative species delimitations, BFD* infers the respective species trees and compares them using Bayes factors [19]. Given the large computational time involved, we ran separate analyses for the two subclades, *A. alpina/A. vesulensis* and *A. pubescens/A. saussurei/A. delphinensis*, but we included all individuals that were assigned to each genomic cluster (*i.e.*, 4 individuals for *A. pubescens*, 5 individuals for *A. vesulensis*, and 9 in *A. saussurei, A. alpina* and *A. delphinensis*). In both cases, *A. helvetica* (represented by 8 individuals only to reduce computational time) was used as outgroup to root species trees, forward and backward mutation rates were arbitrarily set to 1 and the speciation rate for the Yule model of branching was assigned a Gamma prior.

In the first clade, we compared a scenario in which *A. alpina* and *A. vesulensis* would be considered as different species rather than being lumped into a single species (Fig. S10), using MCMC chains of 100,000 steps and 48 steps for path sampling of the marginal likelihood of each scenario. We found decisive support for the first scenario (Bayes Factor = 1,052.42), which leads us to recognize a new species: *A. vesulensis sp. nov.* (see description).

In the other clade, we compared three alternative scenarios (Fig. S10): (i) one in which *A. pubescens, A. saussurei* and *A. delphinensis* would be considered as distinct species, (ii) one in which *A. saussurei* and *A. delphinensis* would be lumped together but *A. pubescens* would remain a distinct species, and (iii) a last one lumping the three lineages into a single species, as assumed by current taxonomy. In this case we used MCMC chains of 250,000 to 300,000 steps and 48 steps for path sampling. Here again, we found decisive support for the scenario in which *A. pubescens, A. saussurei* and *A. delphinensis* would be considered as distinct species, rather than lumping *A. saussurei* and *A. delphinensis* (Bayes Factor = 690.68), or even lumping the three clusters into a single species, as currently done (Bayes Factor = 1,698.15). This leads us to recognize two new species: *A. saussurei sp. nov.* and *A. delphinensis sp. nov.*(see description).

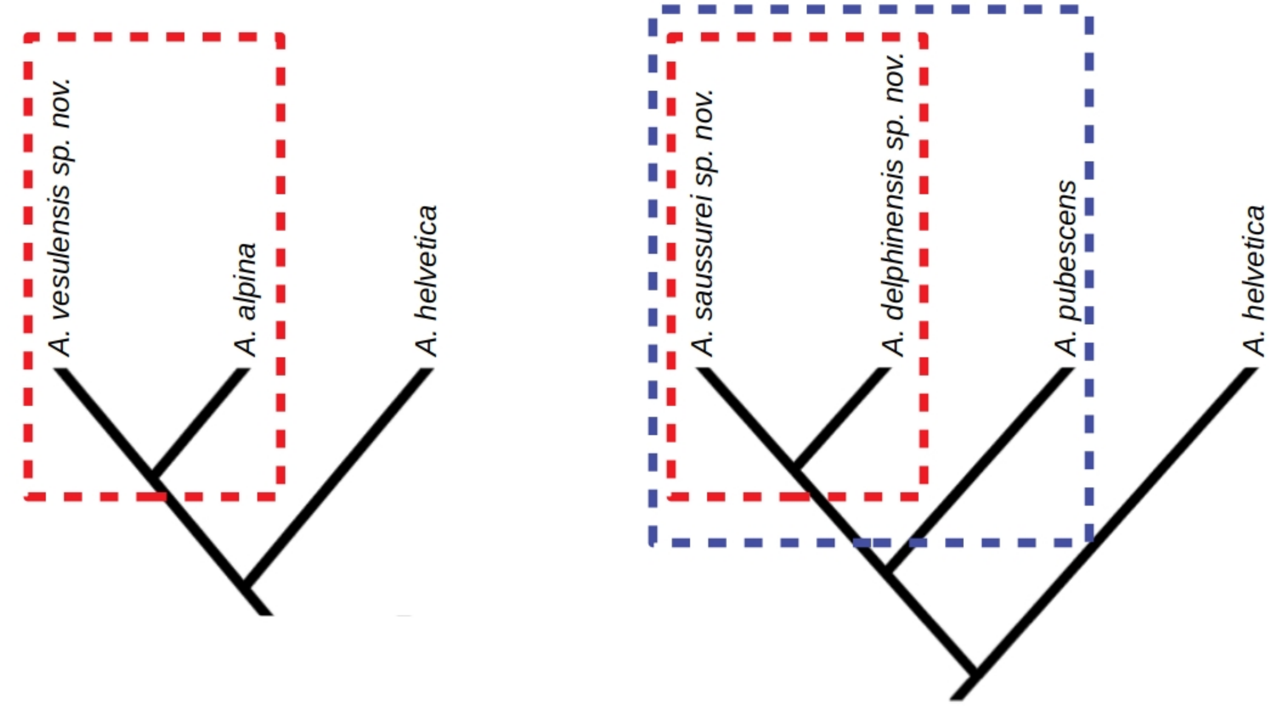


**Fig. S10. Alternative species delimitation scenarios that were compared.** The two sketches depict the two distinct species delimitation analyses that were run. Left: we compared a scenario in with three distinct species to one in which *A. alpina* and *A. vesulensis* would be lumped into a single species (red square). Right: we compared a scenario in which *A. pubescens*, *A. saussurei* and *A. delphinensis* would be considered as distinct species, to one in which *A. saussurei* and *A. delphinensis* would be lumped together (red square) and a last scenario lumping the three lineages into a single species (blue square).

In order to confirm species delimitation results, we visualized genomic distances between the six proposed species, using a genetic PCA as implemented in the *adegenet* R package [21]. This analysis confirmed that all proposed species except for *A. saussurei sp. nov.* and *A. delphinensis sp. nov.* are genetically distinct and not arbitrary portions of a larger genomic cline (Fig. S11).

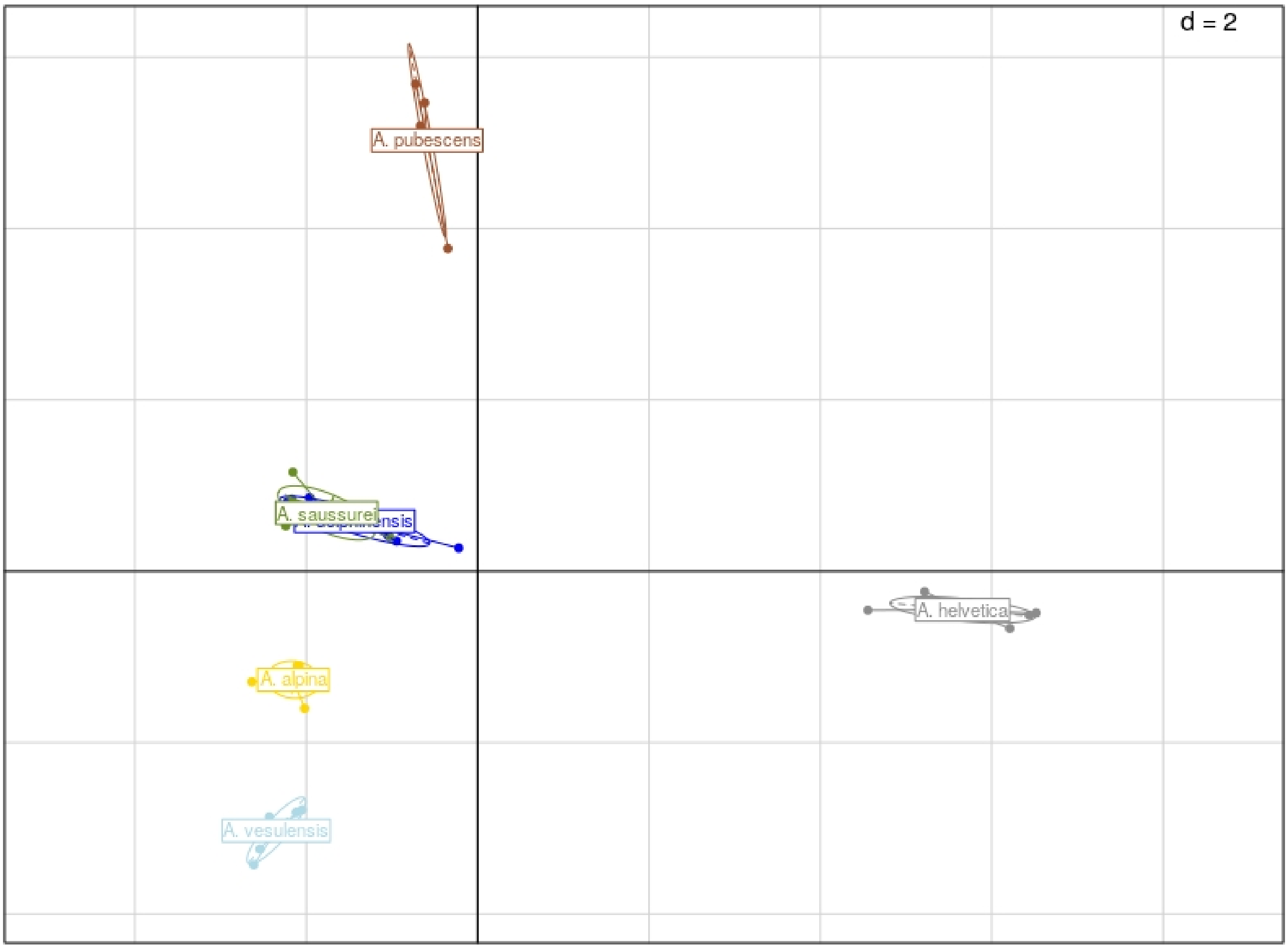


**Fig. S11. Genetic PCA of individuals of /Helvetica.** The graph shows the positions of the 51 individuals from /Helvetica that we sampled along the first two PCA axes, representing 26% and 9% of the total genetic variability of the dataset. Samples have been grouped by the species they belong to and colors match those used in Fig. 3, S5, S6, S8 & S9.

We then looked in more details at the two species that appear to overlap on this first PCA: *A. saussurei sp. nov.* and *A. delphinensis sp. nov.*. Evaluating this species pair is of particular importance since both species have almost identical morphologies and ecologies (see determination key). To do so we ran another PCA including individuals from these two taxa alone, which then confirmed that both species form discrete genomic clusters (Fig. S12).

**Fig. S12.**

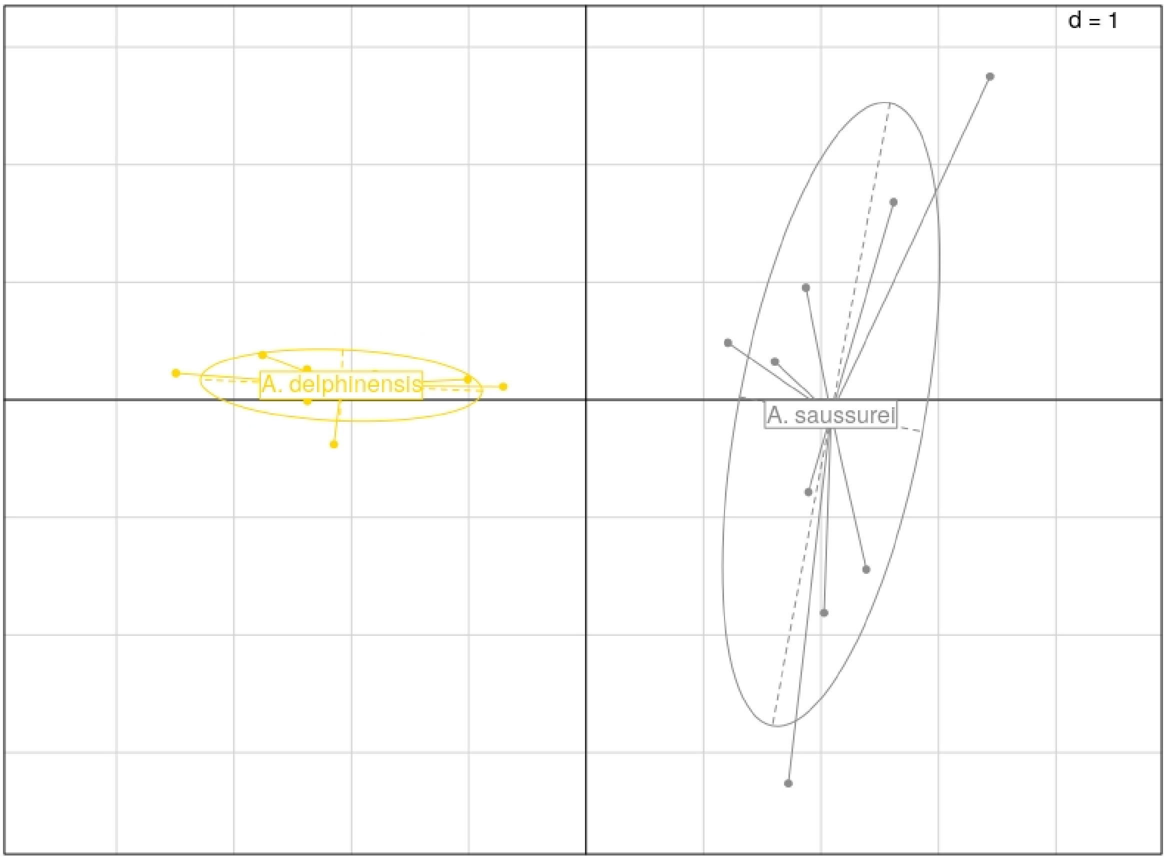


**Genetic PCA of individuals of /Helvetica.** The graph shows the positions of the 18 individuals from *A. delphinensis sp. nov.* and *A. saussurei sp. nov.* that we sampled along the first two PCA axes, representing 32% and 10% of the total genetic variability of the dataset. Samples have been grouped by the species they belong to and colors match those used in Fig. 3, S5, S6, S8, S9 & S11.

Pairwise $F_{ST}$ were then measured between all species pairs using the 'pairwise.fst' function in the R package *hierfstat*[22].Values, averaged over all 381 strictly filtered SNPs, are shown in the table below. $F_{ST}$ between newly described species and their close relatives range from 0.08 to 0.3, in the same order of magnitude as between already described species like *A. pubescens* and *A. alpina* ($F_{ST}$=0.29) or *A. pubescens* and *A. helvetica* ($F_{ST}$=0.17).

| | *A. alpina* | *A. delphinensis* | *A. helvetica* | *A. pubescens* | *A. saussurei* | *A. vesulensis* |
|---|---|---|---|---|---|---|
| *A. alpina* | - | - | - | - | - | - |
| *A. delphinensis* | 0.18 | - | - | - | - | - |
| *A. helvetica* | 0.33 | 0.39 | - | - | - | - |
| *A. pubescens* | 0.12 | 0.13 | 0.17 | - | - | - |
| *A. saussurei* | 0.08 | 0.17 | 0.40 | 0.12 | - | - |
| *A. vesulensis* | 0.30 | 0.15 | 0.22 | 0.12 | 0.19 | - |

Finally, we tested whether the genetic structure that we interpret as evidence of the presence of two distinct species, *A. delphinensis sp. nov.* and *A. saussurei sp. nov.*, could be due to a simple pattern of isolation by distance (hereafter, IBD), as suggested by some authors [23]. To do so we measured the correlation between the genetic and geographic distances separating individuals. Results were similar for the various genetic distances that we used (raw distances, Jukes-Cantor, Kimura or the logDet distance). A mantel correlogram (Fig. S13) showed that genetic distances are strongly and positively correlated to geographic distances at short distances (between 0 and 27.75 km), this correlation remains positive for slightly larger distances (between 27.75 and 55.5 km), but that turns negative for larger distances (over 55.5 km). As this larger distance class corresponds to the spatial distance between the mountain ranges were these two taxa occur, these results show that while IBD is present within *A. saussurei sp. nov.* and *A. delphinensis sp. nov.* separately, it does not exist between them. This two taxa thus cannot be considered as arbitrary ends of a single genetic continuum but rather  represent well delimited genetic lineages.

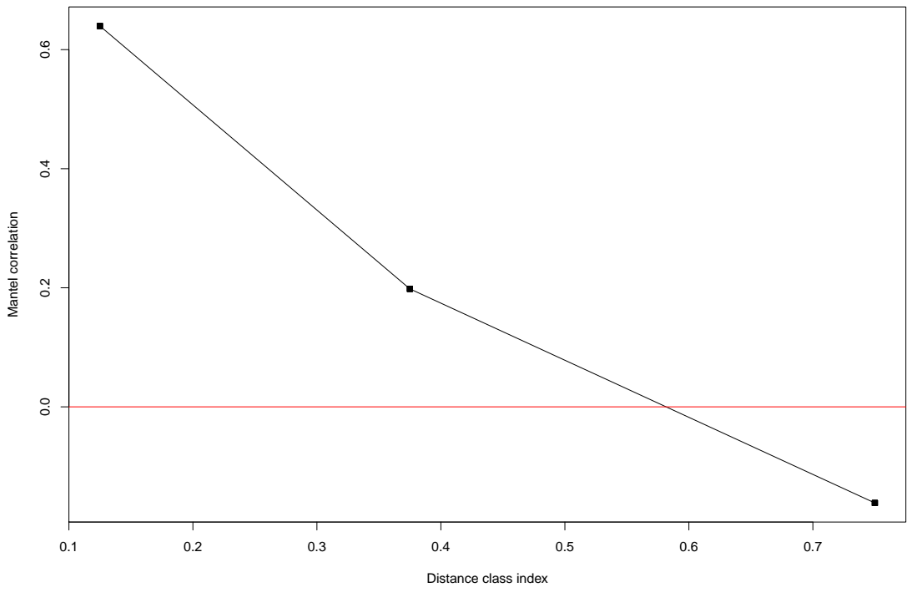


**Fig. S13. Correlation between genetic and geographic distances between individuals of *AA. saussurei* and *delphinensis*.** Genetic distances were measured over full concatenated ddRAD sequences of 276,745bp, using the Jukes-Cantor model. The three points represent the three following geographic distance classes : 0-27.75 km, 27.75-55.5 km and 55.5-150 km. All three correlation coefficients are significantly different from 0 (represented by the red line), as estimated using 999 permutations in the Mantel test.

*3.4. Phylogenetic relationships between newly described species*

Once we had confirmed that these six taxa deserved species status, we inferred their phylogenetic relationships using species tree inference from our strict selection of 381 unlinked SNPs. Here again, we relied on SNAPP to directly estimate a species tree from unlinked biallelic markers without explicitly reconstructing gene trees [20]. In this analysis each of the six species was represented by all individuals for which we had genetic data (51 individuals in total). We ran four independent chains of 500,000 steps, which we then combined after removing the first 10% steps of each, having verified that this led to effective sample sizes > 100 for all parameters of the model. From this combined chain we generated a maximum clade credibility tree with median heights for the nodes, in arbitrary coalescent units. The species tree was then time-calibrated using the crown age of /Helvetica derived from the dated phylogeny of *Androsace* sect. *Aretia* obtained above, *i.e.* 4.66 Myrs.

The topology of this tree was strongly supported: except for the most basal divergence between *A. helvetica* and the rest of the clade, which received only 0.46 posterior support, all other nodes received 1.00 posterior support (Fig. S14). Time calibration suggested that the three newly described species are extremely young: they all probably originated in the last million years (Fig. S14).

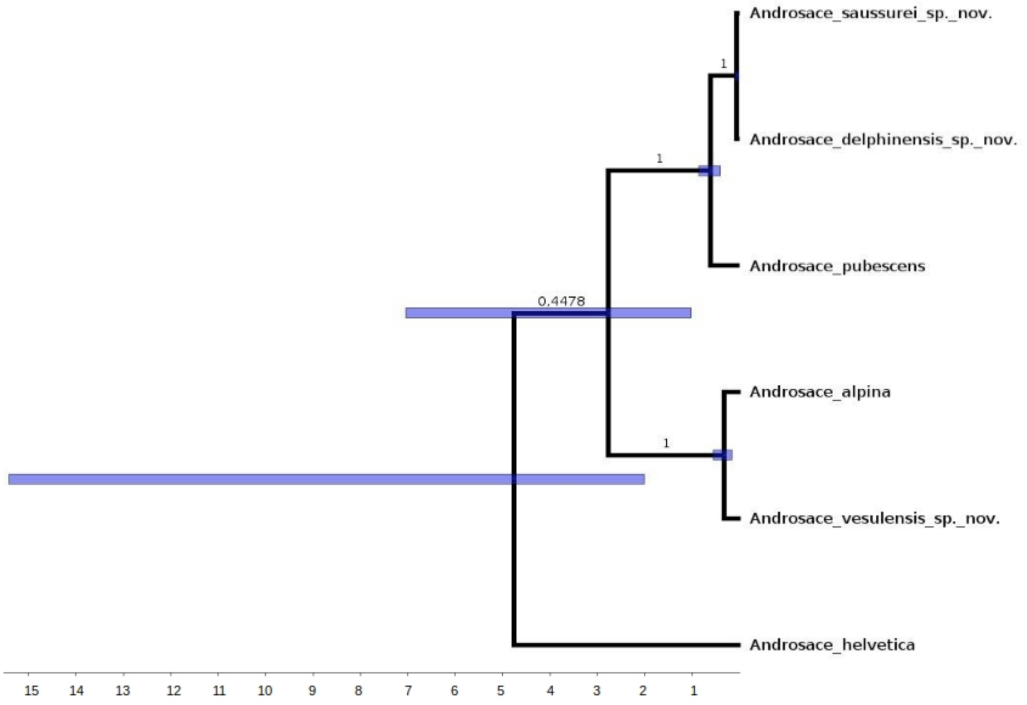


**Fig. S14. Time calibrated species tree of /Helvetica.** Time is shown on the x-axis in million years, with blue bars at node showing the 95% high-probability-density for node ages. Posterior support is shown above branches.

**4. Geographic distribution of species from the /Helvetica clade**

**Fig.**

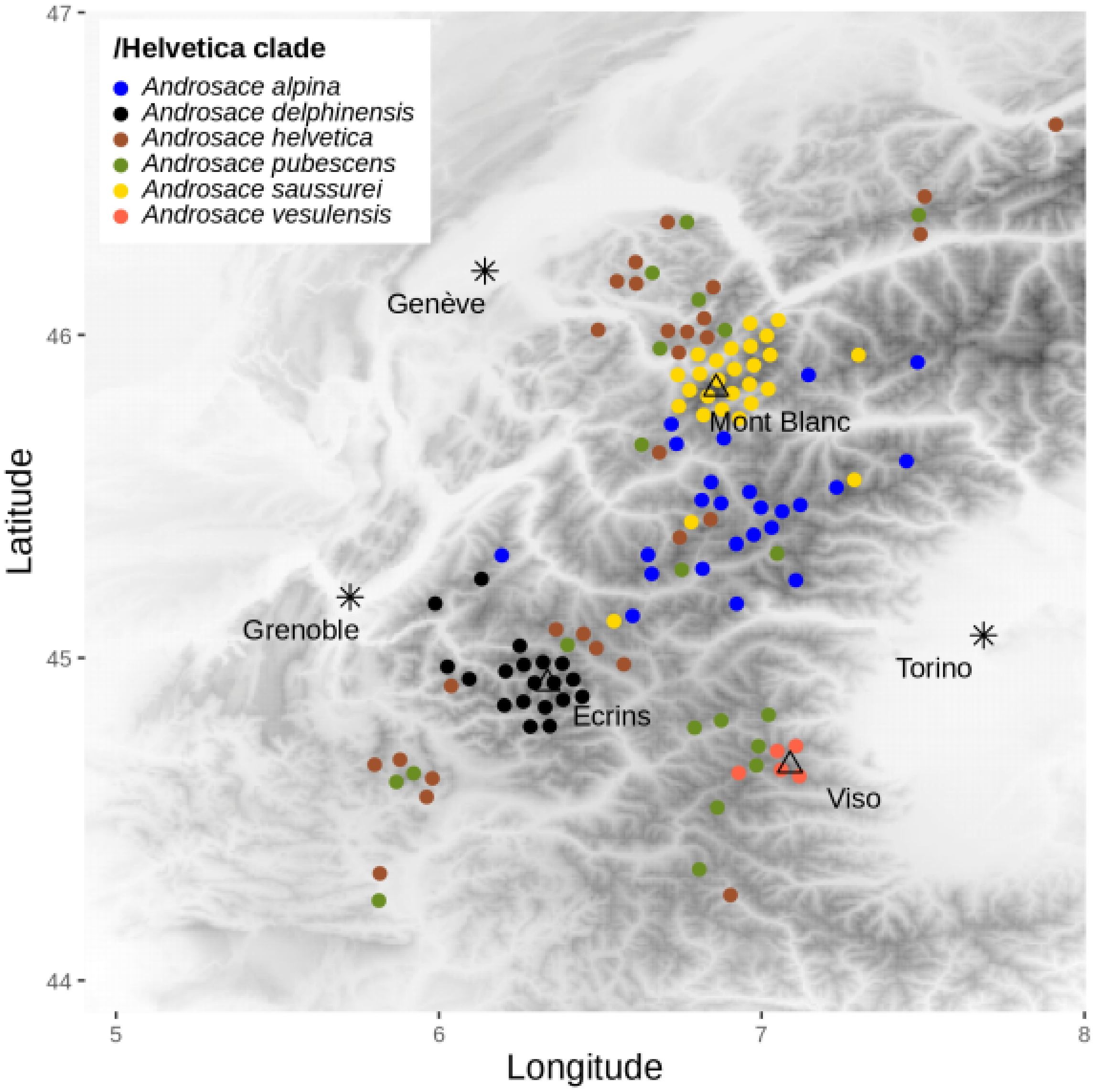


**S15. Updated chorology of the six species from /Helvetica**, based on field collections performed by the authors on this study and on georeferenced herbarium collections checked during the "Herbonautes" citizen science project. The background map shows elevation in shades of grey, main cities are shows with stars and summits with triangles.

**5. Determination key**

1. Flowers white.........................................................................................................................2

1’. Flowers purple, or white to purplish (including buds) on the same cushion ................................7

2. Hairs of leaves strictly simple. Plants growing on limestone......................................................3

2’. At least some hairs on leafs bi-furcated (even with only a small branch at the top). Plants growing on siliceous substrates or ophiolite...............................................................................4

3. Cushions compact and (hemi-)spherical. Leaves short, imbricated, densely hairy. Rosettes dense, forming short and tight columns..................................................................***A. helvetica***

3’. Cushions creeping. Leaves lanceolate, sparsely hairy. Rosettes loose, not forming columns......... ..............................................................................................................***A. pubescens***

4. All leaf hairs clearly deer-antler-shaped..............................................................................5

4’. Leaf hairs simple and bifurcated, tri-furcated mainly on pedicels................................................6

5. Hairs of two types : the shortest up to 0.1 mm long, the biggest up to 0.2 mm long. Plant forming a cushion, more or less loose. Plant of Monte Viso (Italy) and surroudings..........***A. vesulensis***

5’. Hairs of only one type, 0.1 mm long. Plant forming a creeping cushion. Plant of siliceous substrates (Alps)...............................................................................................***A. alpina***

6. Rosettes dense. Leaves often with reddish tips. Plant of the Mont Blanc range..........***A. saussurei***

6’. Rosettes loose. Leaves rarely with reddish tips. Plant of the Ecrins range.......... ***A. delphinensis***

7. Hairs 0.1 mm long (on leaves, pedicels, calyx), strictly deer-antler-shaped. Plant forming a creeping cushion................................................................................................***A. alpina***

7’. Hairs more than 0.2 mm long, simple and bifurcated on leaves, mainly tri-furcated on pedicels. Plant forming a denser cushion ***A. saussurei***

#### References

1. Peterson, B. K., Weber, J. N., Kay, E. H., Fisher, H. S. & Hoekstra, H. E. Double Digest RADseq: An Inexpensive Method for De Novo SNP Discovery and Genotyping in Model and Non-Model Species. *PLoS One* **7**, e37135 (2012).
2. Nowak, M. D. *et al.* The draft genome of Primula veris yields insights into the molecular basis of heterostyly. *Genome Biol.* **16**, 12 (2015).
3. Boucher, F. C., Casazza, G., Szövényi, P. & Conti, E. Sequence capture using RAD probes clarifies phylogenetic relationships and species boundaries in Primula sect. Auricula. *Mol. Phylogenet. Evol.* **104**, 60–72 (2016).
4. Boucher, F. C., Zimmermann, N. E. & Conti, E. Allopatric speciation with little niche divergence is common among alpine Primulaceae. *J. Biogeogr.* **43**, 591–602 (2016).
5. Stamatakis, A. RAxML version 8: A tool for phylogenetic analysis and post-analysis of large phylogenies. *Bioinformatics* **30**, 1312–1313 (2014).
6. Nguyen, L.-T., Schmidt, H. A., von Haeseler, A. & Minh, B. Q. IQ-TREE: a fast and effective stochastic algorithm for estimating maximum-likelihood phylogenies. *Mol. Biol. Evol.* **32**, 268–274 (2015).
7. Rannala, B. & Yang, Z. Bayes estimation of species divergence times and ancestral population sizes using DNA sequences from multiple loci. *Genetics* **164**, 1645–1656 (2003).
8. Chifman, J. & Kubatko, L. Quartet Inference from SNP Data Under the Coalescent Model. *Bioinformatics* **30**, 3317–3324 (2014).
9. Schneeweiss, G., Schönswetter, P., Kelso, S. & Niklfeld, H. Complex biogeographic patterns in Androsace (Primulaceae) and related genera: evidence from phylogenetic analyses of nuclear internal transcribed spacer and plastid trnL-F sequences. *Syst. Biol.* **53**, 856–876 (2004).
10. Dixon, C. J., Schönswetter, P. & Schneeweiss, G. M. Traces of ancient range shifts in a mountain plant group (Androsace halleri complex, Primulaceae). *Mol. Ecol.* **16**, 3890–3901 (2007).
11. Boucher, F. C. *et al.* Reconstructing the origins of high-alpine niches and cushion life form in the genus Androsace s.l. (Primulaceae). *Evolution (N. Y).* **66**, 1255–1268 (2012).
12. Sanderson, M. J. Estimating Absolute Rates of Molecular Evolution and Divergence Times: A Penalized Likelihood Approach. *Mol. Biol. Evol.* **19**, 101–109 (2002).
13. Paradis, E. Molecular dating of phylogenies by likelihood methods: A comparison of models and a new information criterion. *Mol. Phylogenet. Evol.* **67**, 436–444 (2013).
14. Forest, F. Calibrating the tree of life: Fossils, molecules and evolutionary timescales. *Ann. Bot.* **104**, 789–794 (2009).

15. Janzen, D. H. *et al.* Nuclear genomes distinguish cryptic species suggested by their DNA barcodes and ecology. (2017). doi:10.1073/pnas.1621504114

16. Devitt, T. J., Wright, A. M., Cannatella, D. C. & Hillis, D. M. Species delimitation in endangered groundwater salamanders : Implications for aquifer management and biodiversity conservation. 1–10 (2018). doi:10.1073/pnas.1815014116

17. Frichot, E., Mathieu, F., Trouillon, T., Bouchard, G. & François, O. Fast and Efficient Estimation of Individual Ancestry Coefficients. *Genetics* **196**, 973 LP – 983 (2014).

18. Durand, E. Y., Patterson, N., Reich, D. & Slatkin, M. Testing for ancient admixture between closely related populations. *Mol. Biol. Evol.* **28**, 2239–2252 (2011).

19. Leaché, A. D., Fujita, M. K., Minin, V. N. & Bouckaert, R. R. Species delimitation using genome-wide SNP data. *Syst. Biol.* **63**, 534–542 (2014).

20. Bryant, D., Bouckaert, R., Felsenstein, J., Rosenberg, N. A. & RoyChoudhury, A. Inferring species trees directly from biallelic genetic markers: bypassing gene trees in a full coalescent analysis. *Mol. Biol. Evol.* **29**, 1917–1932 (2012).

21. Jombart, T. & Ahmed, I. adegenet 1.3-1: new tools for the analysis of genome-wide SNP data. *Bioinformatics* **27**, 3070–3071 (2011).

22. Goudet, J. hierfstat, a package for r to compute and test hierarchical F-statistics. *Mol. Ecol. Notes* **5**, 184–186 (2005).

23. Chambers, E. A. & Hillis, D. M. The Multispecies Coalescent Over-Splits Species in the Case of Geographically Widespread Taxa. *Syst. Biol.* **69**, 184–193 (2019).
